# *β*2 integrins impose a mechanical checkpoint on macrophage phagocytosis

**DOI:** 10.1101/2024.02.20.580845

**Authors:** Alexander H. Settle, Benjamin Y. Winer, Miguel M. de Jesus, Lauren Seeman, Zhaoquan Wang, Eric Chan, Yevgeniy Romin, Zhuoning Li, Matthew M. Miele, Ronald C. Hendrickson, Daan Vorselen, Justin S. A. Perry, Morgan Huse

**Affiliations:** Louis V. Gerstner, Jr., Graduate School of Biomedical Sciences, Memorial Sloan Kettering Cancer Center, New York, NY USA; Immunology Program, Memorial Sloan Kettering Cancer Center, New York, NY USA; Immunology and Molecular Pathogenesis Program, Weill Cornell Medicine Graduate School of Medical Sciences, New York, NY USA; Molecular Cytology Core Facility, Memorial Sloan Kettering Cancer Center, New York, NY USA; Proteomics Core Facility, Memorial Sloan Kettering Cancer Center, New York, NY USA; Molecular Pharmacology Program, Memorial Sloan Kettering Cancer Center, New York, NY USA; Cell Biology and Immunology, Wageningen University Research, Wageningen, the Netherlands; Department of Biochemistry and Molecular Biology, University of Miami School of Medicine, Miami, FL

## Abstract

Phagocytosis is an intensely physical process that depends on the mechanical properties of both the phagocytic cell and its chosen target. Here, we employed differentially deformable hydrogel microparticles to examine the role of cargo rigidity in the regulation of phagocytosis by macrophages. Whereas stiff cargos elicited canonical phagocytic cup formation and rapid engulfment, soft cargos induced an architecturally distinct response, characterized by filamentous actin protrusions at the center of the contact site, slower cup advancement, and frequent phagocytic stalling. Using phosphoproteomics, we identified *β*2 integrins and their downstream effectors as critical mediators of this mechanically regulated phagocytic switch. Indeed, comparison of wild type and *β*2 integrin deficient macrophages indicated that integrin signaling acts as a mechanical checkpoint by shaping filamentous actin to enable distinct phagocytic engulfment strategies. Collectively, these results illuminate the molecular logic of leukocyte mechanosensing and reveal potential avenues for modulating phagocyte function in immunotherapeutic contexts.

## Introduction

Macrophages maintain homeostasis in multicellular organisms by phagocytosing microbes, cell corpses, and biomolecular debris. They also play a central role in emerging immunotherapies against infectious diseases and cancer, which has spawned intense interest in the mechanisms that control their activity (Accarias, Sanchez et al. 2020, Klichinsky, Ruella et al. 2020, Mantovani, Allavena et al. 2022, Sly and McKay 2022). Macrophages and other professional phagocytes, including neutrophils and dendritic cells, consume cargo that bears the molecular indices of immune targeting (antibodies and complement), microbial origin (e.g. lipopolysaccharide), or cellular distress (e.g. phosphatidylserine (PtdS)). Recognition of these ligands is mediated by phagocytic receptors specific for each ligand class: Fc receptors and *β*2 integrins bind to opsonizing antibodies and the iC3b subunit of complement, respectively, TLR4 binds to lipopolysaccharide, and a structurally diverse group of cell surface proteins recognizes PtdS. Receptor engagement induces dramatic reorganization of filamentous (F)-actin at the interface, which shapes the overlying plasma membrane into a phagocytic cup that engulfs the cargo, ultimately leading to its internalization in an acidifying phagosome.

Phagocytic cargo varies widely in its chemical composition, architecture, and mechanical properties, and phagocytes have developed multiple uptake strategies to accommodate this diversity (Swanson and Baer 1995, Jaumouillé and Waterman 2020). The morphological features of phagocytosis have traditionally been linked to specific biochemical modes of target engagement. Thus, antibody-induced engulfment is thought to occur via a “reaching” mechanism in which F-actin rich pseudopods promote the sequential engagement (zippering) of Fc receptors, enabling the phagocyte to surround and then consume its target. Conversely, complement dependent uptake has been proposed to proceed by a “sinking” mechanism in which cargo is pulled into the phagocyte with minimal formation of membrane pseudopods. These distinctions have been challenged, however, by studies highlighting architectural similarities between Fc- and complement-mediated phagocytosis, in particular the importance of protrusive F-actin for both processes (Jaumouille, Cartagena-Rivera et al. 2019, Walbaum, Ambrosy et al. 2021). More recent work has also revealed an alternative internalization response in which cargo is fragmented before consumption (Velmurugan, Challa et al. 2016, Vorselen, Kamber et al. 2022, Zhao, Zhang et al. 2022). This “nibbling” process has been proposed to facilitate collaborative phagocytosis, which may be essential for the consumption of large cargos (Dooling, Andrechak et al. 2023). Deciphering how and why distinct uptake mechanisms are applied will require a comprehensive understanding of how macrophages sense and classify their cargo.

In considering this question, it is important to bear in mind that phagocytic cargos vary not only biochemically but also mechanically, ranging from hard, elongated bacterial cells to irregularly shaped and highly deformable apoptotic corpses (Eaton, Fernandes et al. 2008, Bufi, Saitakis et al. 2015, Planade, Belbahri et al. 2019). The implications of this biophysical diversity for phagocytosis, which is itself and intensely physical process (Jaumouillé and Waterman 2020, Vorselen, Labitigan et al. 2020), are likely profound. Within the phagocytic cup, macrophages apply nanonewton scale protrusive forces to palpate the target surface (Vorselen, Barger et al. 2021) potentially providing an avenue for cargo mechanosensing. Among the physical properties that could be detected in this manner, rigidity is particularly interesting as a candidate regulator of phagocytosis because it varies substantially among cargos. Polystyrene beads and red blood cells, for instance, differ by six orders of magnitude in their Young’s Modulus (Bufi, Saitakis et al. 2015). Prior studies have explored the effects of cargo rigidity on phagocytosis using hydrogel microparticles of differential stiffness as well as red blood cell targets treated with fixatives like glutaraldehyde. In general, increased rigidity was found to enhance uptake, indicating that phagocytes are indeed mechanosensitive and that they prefer stiffer cargo (Beningo and Wang 2002, Sosale, Rouhiparkouhi et al. 2015, Jaumouille, Cartagena-Rivera et al. 2019, Vorselen, Wang et al. 2020). The cellular and mechanical bases that underlie this preference, however, remain largely unknown.

In the present study, we investigated the specific effects of cargo stiffness on macrophage phagocytosis. Central to our efforts was the application of microfabrication methodology to generate phagocytic targets of defined size, rigidity, and surface ligand composition (Vorselen, Wang et al. 2020). Our results demonstrate that cargo mechanosensing is a general property of macrophages that is independent of cell lineage and mode of target recognition. We also found that increasing cargo stiffness is associated with stereotyped changes in both the speed and the morphology of phagocytic contacts, indicating that macrophages apply distinct phagocytic programs depending on the mechanical properties of their targets. Finally, we showed that *β*2 integrins act as a mechanical check-point that pairs cargo mechanics to the mechanism of uptake.

## Results

### Cargo rigidity promotes phagocytic uptake

To isolate the role of cargo stiffness on phagocytosis, we employed a recently developed method for generating deformable polyacrylamide acrylic acid micro (DAAM) particles of differential rigidity (Vorselen, Wang et al. 2020) (Fig 1A). For this study, we prepared four sets of particles, all 9-11 µm in diameter (Fig 1B, Supp Fig 1A), with stiffnesses (Young’s moduli) ranging from 0.6 kPa to 18 kPa (Fig 1C), encompassing the range between leukocytes (low rigidity) and yeast (high rigidity). Particles were loaded with phagocytic ligands (IgG or PtdS) via a streptavidin-biotin linkage in order to induce macrophage uptake. Using confocal microscopy, we confirmed that a coating ratio of 10 pmol IgG per million particles (pmol/M) affords particles of identical coating density (Supp Fig 1B,C). Unless otherwise stated, this coating density was used for all phagocytosis assays.

**Fig. 1.**
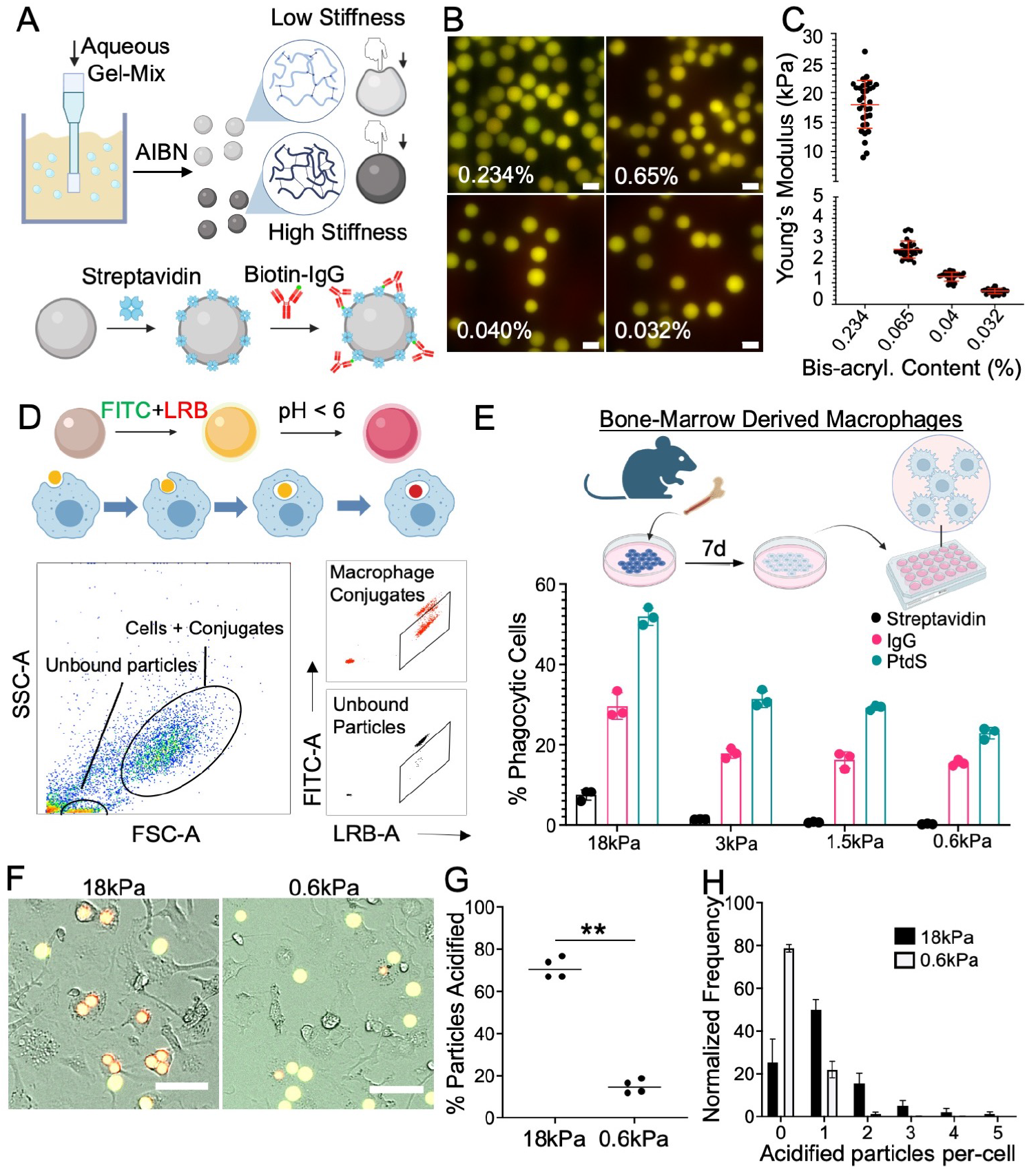
A hydrogel microparticle system to interrogate cargo mechanosensing. A) Schematic diagram for DAAM particle synthesis (see Materials and Methods). Particle rigidity depends on the ratio of cross-linker to monomer. AIBN: Azobisisobutyronitrile. B) Representative images of DAAM particles, prepared using the indicated bisacrylamide contents and conjugated to LRB and FITC. Scale bars = 10 µm. C) Atomic force microscopy measurements of stiffness (Young’s Modulus) for particles prepared using the indicated bis-acrylamide contents. D-E) BMDMs were challenged with DAAM particles coated with different phagocytic ligands and particle uptake quantified by flow cytometry. D) Top, schematic diagram for LRB/FITC dependent tracking of phagocytosis. Bottom, representative gating for flow cytometry-based quantification. E) Uptake efficiency of DAAM particles coated with the indicated phagocytic ligands is graphed against particle stiffness. Columns and error bars denote mean and standard deviation (SD), respectively. n = 3 biological replicates. F-H) BMDMs were imaged together with IgG-coated DAAM particles. F) Representative brightfield and epifluorescence images of BMDMs phagocytosing DAAM particles of the indicated stiffness. Phagocytosed particles turn red, while unengulfed particles are yellow. Scale bars = 50 µm. G) Phagocytosis quantified as the fraction of particles acidified per field of view after 2 h. Each point represents the mean of four fields of view in one technical replicate. Horizontal lines in each column denote the overall mean. ^**^: p<0.01, two-tailed Student’s t-test. H) Histogram showing the number of 18 kPa and 0.6kPa particles phagocytosed per cell after 2 h. Error bars denote SD, calculated from 4 technical replicates.

For most of our experiments, DAAM particles were conjugated to FITC, a pH sensitive dye that fluoresces green, and Lissamine Rhodamine B (LRB), a pH insensitive red fluorophore. Uptake into an acidic phagolysosome quenches FITC fluorescence, enabling the identification of phagocytic macrophages by flow cytometry as LRB+FITClo cells (Fig 1D, Supp Fig 1D-E). In this manner, we quantified the phagocytic activity of primary murine bone marrow derived macrophages (BMDMs) as a function of DAAM particle stiffness. These experiments revealed a distinct preference of macrophages for stiffer cargos; uptake increased monotonically with particle rigidity for both IgG-dependent and PtdS-dependent phagocytosis (Fig 1E). We observed a similar pattern of results over a range of ligand coating densities (Supp Fig 1F), indicating that mechanosensing occurs independently of target quality. We also performed live widefield microscopy of BMDMs co-incubated with stiff (18 kPa) or soft (0.6 kPa) DAAM particles bearing IgG or PtdS (Supp Fig 2A). Automated analysis of the imaging data indicated that the stiff cargo was more effectively engulfed on both a population level and a per-cell basis, with 24 % of macrophages taking up two or more particles (Fig. 1F-H, Supp Fig 2B-E). High stiffness also augmented the uptake of particles coated with streptavidin alone, BSA, or no protein at all, suggesting that this effect does not require a specific phagocytic receptor-ligand interaction (Supp Fig 2F). Collectively, these results demonstrate that stiffness promotes the phagocytosis of chemically diverse cargos, which is consistent with prior reports (Beningo and Wang 2002, Sosale, Rouhiparkouhi et al. 2015, Jaumouille, Cartagena-Rivera et al. 2019, Vorselen, Wang et al. 2020).

Next, we assessed the generality of this behavior by measuring DAAM particle phagocytosis by different macrophage populations and in different contexts. Murine macrophages derived from HoxB8-ER-immortalized progenitor cells (Wang, Calvo et al. 2006, Accarias, Sanchez et al. 2020) displayed a similar preference for stiffer cargo in both IgG and PtdS dependent phagocytosis assays (Fig 2A). We observed analogous results using human macrophages differentiated from induced pluripotent stem cells (hiPSCs) (Fig 2B), indicating that cargo mechanosensing is not a species-specific phenomenon. We also examined microglia, the professional phagocytes of the central nervous system, which reside and operate in an environment that is markedly softer than that of other macrophage subsets. Murine microglia consumed substantially more stiff (18 kPa) than soft (0.6 kPa) DAAM particles (Fig. 2C), strongly suggesting that this form of mechanosensing applies to most, if not all, macrophages. To assess whether the preference for stiff cargo also applies in vivo, we injected IgG-coated particles of differential rigidity into the peritonea of adult mice and measured phagocytosis by peritoneal macrophages after 2 hours (Fig 2D, Supp Fig 3). In vivo uptake of stiff particles was robust, whereas uptake of soft particles was barely detectable (Fig. 2E). Importantly, macrophage recruitment to the peritoneum was comparable in both injection regimes (Fig 2F), indicating that the observed preference for stiff particles did not result from differences in macrophage-particle encounter frequency. Together, these results suggest that cargo mechanosensing is a general phenomenon that applies to a wide range of macrophage cell types and environments.

**Fig. 2.**
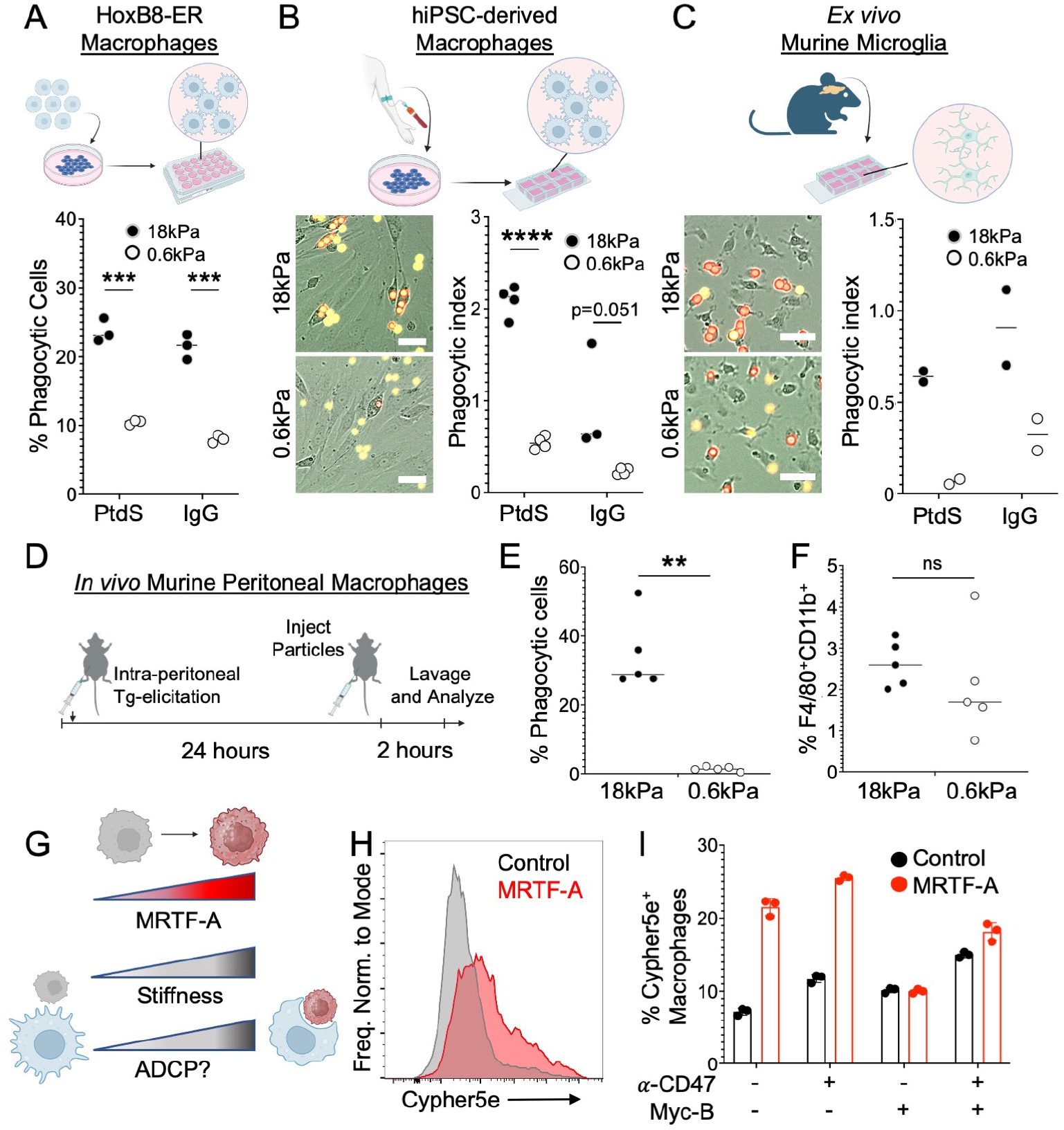
High cargo rigidity promotes phagocytic uptake. A) Phagocytic efficiency of HoxB8-ER derived macrophages challenged with IgG- and PtdS-coated DAAM particles of the indicated rigidities, quantified by flow cytometry. ^***^: p<0.001, two-tailed Student’s t-test. n = 3 replicate macrophage differentiations. B-C) hiPSC-derived macrophages (B) and murine microglia (C) were mixed with IgG- and PtdS-coated DAAM particles of the indicated rigidities, and phagocytosis monitored by videomicroscopy. Representative images of phagocytic uptake are shown to the left, with phagocytic index quantified on the right. Phagocytic index = number of particles phagocytosed per macrophage. ^****^: p<0.0001, two-tailed Student’s t-test. In B, n = 4 replicate macrophage differentiations, while in C, n = 2 biological replicates (summary of four technical replicates per biological replicate). D-F) DAAM particles of differential stiffness were injected into the mouse peritoneum to measure the phagocytic activity of elicited macrophages. D) Schematic of macrophage elicitation and DAAM particle injection protocol (see Materials and Methods). E) Phagocytic efficiency (LRB+FITClo / all CD11b+F4/80+ cells) for particles of the indicated rigidities. n = 5 mice per condition, ^**^: p<0.005, two-tailed Student’s t test. F) Macrophage recruitment to the peritoneum, quantified as total F480+CD11b+ cells over all live cells in the peritoneal lavage. ns: not significant, two-tailed Student’s t test. In A-C and E-F, horizontal lines in columns denote mean values. G-I) BMDMs were challenged with Cypher5e-labeled E0771 cells expressing different levels of MRTF-A. G) MRTF-A is expected to enhance ADCP by increasing cell stiffness. H) Representative flow cytometry histogram showing Cypher5e signal in BMDMs challenged for 2 h with control E0771 cells or E0771 cells overexpressing MRTF-A. I) Phagocytosis quantified as % Cypher5e+ macrophages. Error bars indicated SD. n = 3 replicate macrophage differentiations.

In light of these observations, we hypothesized that mechanically rigidifying cancer cells would promote their phagocytosis by macrophages. Overexpression of myocardin related transcription factors (MRTFs) increases the Young’s Modulus of multiple cell types without substantially altering their surface composition (Tello-Lafoz, Srpan et al. 2021) (Fig 2G). Accordingly, we challenged BMDMs with E0771 breast cancer cells bearing an inducible MRTF-A expression cassette. Phagocytosis was measured in both the presence and the absence of anti-CD47 antibodies, which were used to block phagocytic “don’t eat me” signaling. MRTF-A over-expression markedly increased E0771 uptake in these experiments, independently of anti-CD47 treatment (Fig 2H-I). To determine whether this result depended specifically on the cytoskeletal effects of MRTF-A, target cells were incubated with mycalolide B (MycB), a small molecule toxin that depolymerizes F-actin in a sustained manner (Saito, Watabe et al. 1994). MycB pretreatment completely abrogated the enhanced phagocytosis of MRTF-A overexpressing cells (Fig 2I), strongly suggesting that MRTF-induced rigidification is the critical determinant of this effect. Hence, cargo stiffness promotes the uptake of both live and inanimate phagocytic cargos.

### Phagocytosis of soft cargo is slow and prone to stalling

To better understand the underlying mechanism of cargo mechanosensing, we analyzed phagocytic kinetics in wide-field videos of BMDMs engulfing stiff (18 kPa) and soft (0.6 kPa) DAAM particles (Fig 3 A-B). For each engulfment event, we scored three key steps: (1) contact, defined as the first frame the particle contacts the cell, (2) phagocytic cup initiation, visualized as an inversion in brightfield intensity around the particle, and (3) particle acidification, defined as the frame at which the FITC/LRB ratio reaches its minimum plateau (see Materials and Methods). Whereas stiffness did not affect the time delay between particle contact and cup initiation (Fig 3C, Supp Fig 4A), the interval between initiation and particle acidification was significantly faster dur-ing stiff phagocytosis (Fig 3D, Supp Fig 4B). Given that the maximal rate of soft particle acidification did not vary significantly from that of stiff particles (Supp Fig 4C), we surmised that phagocytic mechanosensing occurs between cup initiation and closure. Guided by these results, we turned to high speed (15 s interval) confocal microscopy to measure the rate of phagocytic cup closure more directly. These experiments utilized HoxB8-ER macrophages expressing f-Tractin-mCherry, a probe for F-actin, which we mixed with fluorescently labeled DAAM particles just prior to imaging. F-actin rapidly accumulated at both stiff particle and soft particle interfaces. On stiff (18 kPa) particles, this accumulation promptly transitioned into a radially symmetric phagocytic cup containing a thick band of leading-edge F-actin, which advanced rapidly (0.12 um/s) around the cargo, leading to full engulfment within 10 minutes of contact formation (Fig 3E-G, Supp Fig 4D, Supp Video 1). Soft (600 Pa) particle contacts were markedly different in character: although macrophages appeared to form phagocytic cups on the target, most (10/15) failed to complete engulfment within 30 minutes, with little to no advancement of the phagocytic cup during that time (Fig 3E-G, Supp Fig 4D-E, Supp Videos 2-3). This behavior, which we defined as “stalled phagocytosis”, was not observed in contacts with stiff particles (0/13) (Fig 3G). Even in the minority of cases where soft particle engulfment did complete, phagocytic cup advancement was approximately 3-fold slower than the rate observed with stiff particles (Fig 3F).

**Fig. 3.**
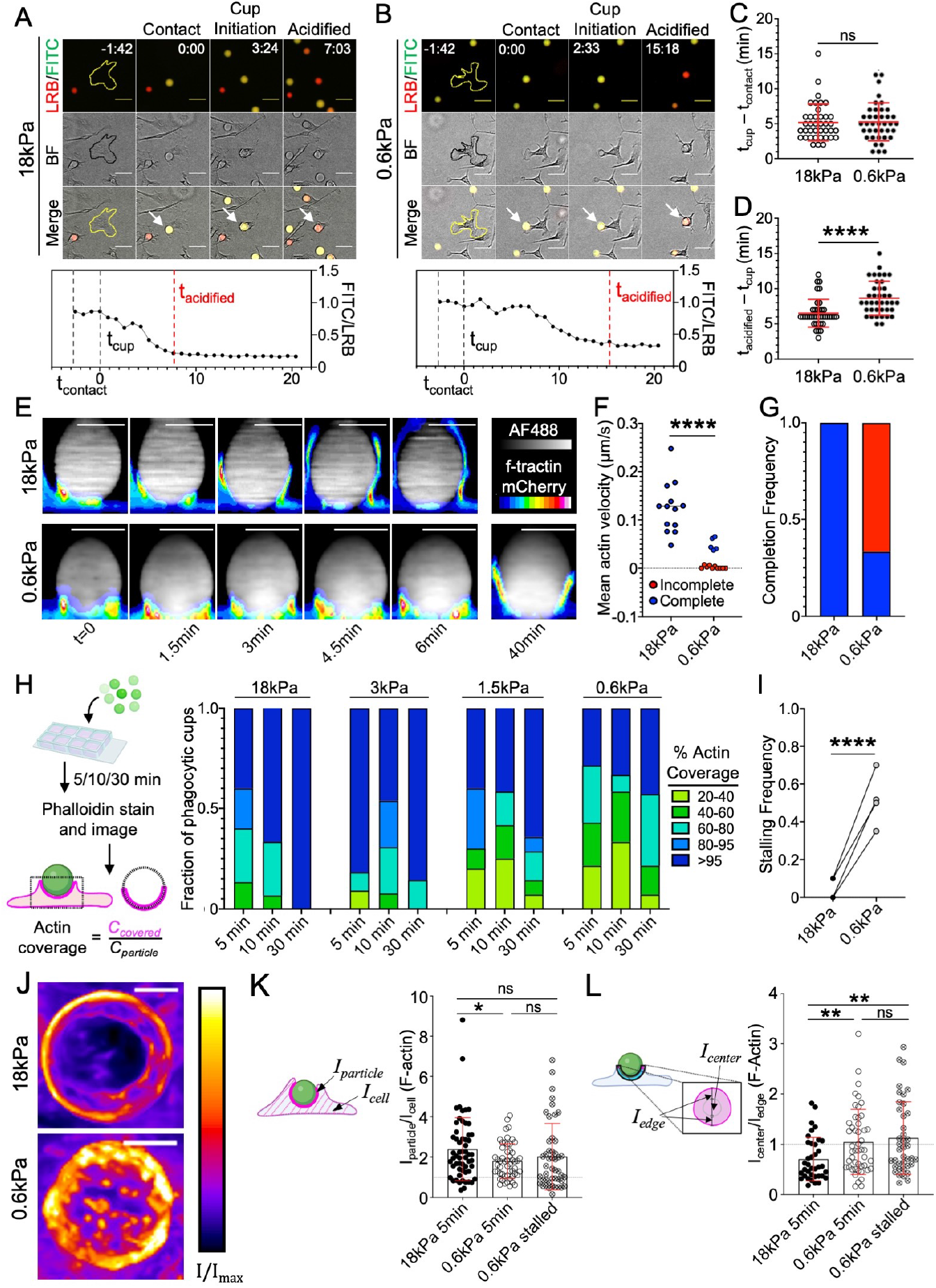
Phagocytosis of soft targets is slow and prone to stalling. A-C) BMDMs were challenged with 18 kPa or 0.6 kPa IgG-coated DAAM particles and imaged by widefield videomicroscopy. A-B) Above, time-lapse montages showing representative 18 kPa (A) and 0.6 kPa (B) uptake events. Fluorescence (LRB/FITC) and brightfield (BF) channels are shown individually and in merge. Particles undergoing phagocytosis are indicated by white arrows in the merged images. The BMDM of interest is outlined in yellow/black in the first image of each time-lapse. Time is indicated as M:SS in the top right of the fluorescence images, with time of first contact defined as t=0. Scale bars = 30 µm. C) Recognition time, defined as time difference between cup formation and first contact, n=39 cells. ns: not significant, two-tailed Welch’s t-test. D) Total acidification time, defined as the difference between particle acidification and cup formation, n = 39 cells, ^****^: p<0.0001, two-tailed Welch’s t-test. In C and D, mean values and error bars (denoting SD) are indicated in red. E-G) HoxB8-ER macrophages expressing f-tractin-mCherry were challenged with stiff (18 kPa) or soft (0.6 kPa) IgG-coated DAAM particles and imaged by spinning disk confocal microscopy. E) Time-lapse montages of representative phagocytic events, visualized as sideviews of 3D reconstructions. f-tractin is depicted in pseudocolor, with warmer colors indicating higher fluorescence, and the particle is shown in gray. A 40-min time point from the same 0.6 kPa particle conjugate is included for reference. Scale bars = 10 µm. F) Mean phagocytic cup velocity (see Materials and Methods and Supp Fig 4D-E) on stiff and soft DAAM particles. Blue and red points denote cups that completed or failed to complete within 30 min, respectively. Statistical significance (p<0.0001) includes all events, p = 0.0002 using only completed events, two-tailed Welch’s t-test. G) Fraction of events that completed (blue) or did not complete (red) within 30 min. Completion defined as actin covering >95% of the particle at any time and persisting until the end of capture. H-I) BMDMs were challenged with IgG-coated DAAM particles of different rigidities for various times, and then fixed and stained with phalloidin (F-actin). H) Left, Schematic for quantification of % actin coverage on fixed phagocytic cups. Right, F-actin coverage for various particle rigidities, graphed against time. Values are sorted into five bins, with colors denoting the fraction of events falling within each bin. n=15 events per time/stiffness condition (180 total events) in one representative experiment. I) Quantification of stalling frequency, defined as the fraction of events that are 20-95 % complete after 30 min co-incubation with BMDMs. ^****^: p<0.0001, two-tailed paired t-test, n=4 biological replicates (one mouse per replicate). J) Representative en-face maximum z-projections of partial phagocytic cups formed on 18 kPa and 0.6 kPa DAAM particles, taken from live imaging experiments. K-L) F-actin accumulation (K), defined as the ratio of phalloidin stain at the particle interface divided by total phalloidin stain in the cell, and F-actin clearance ratio (L), which compares F-actin signal at the periphery and the center of the phagocytic cup (see Materials and Methods), determined in phagocytic cups formed with 18 kPa (5 minutes) and 0.6 kPa (5 or 30 minutes) DAAM particles. In K and L, error bars denote SD. ^*^: p < 0.05, ^**^: p < 0.01, two-tailed Welch’s t-test. n=37, 48, 40 cells for 18 kPa, 0.6 kPa 5min, and 0.6kPa 30min, respectively.

To confirm that phagocytic stalling is indeed a function of target rigidity, we imaged conjugates of BMDMs and IgG-coated DAAM particles, which had been fixed and stained for F-actin at various times after mixing (Fig. 3H). At early timepoints (5 and 10 minutes), samples containing stiff (18 kPa) particles exhibited both partially and fully internalized beads, indicative of ongoing and completed phagocytosis, respectively (Fig 3H). After 30 minutes, all observed 18 kPa particles were fully internalized, implying that all phagocytic cups had fully enclosed their cargo by this time point. By contrast, samples containing softer DAAM particles (3 kPa, 1.5 kPa, 0.6k Pa) exhibited a substantial degree of incomplete engulfment at 30 minutes, consistent with the formation of stalled phagocytic cups (Fig. 3H-I). Notably, the fraction of stalled events at 30 minutes increased monotonically with decreasing particle rigidity, strongly suggesting that the likelihood to stall directly reflects target deformability. Hence, at the level of individual cells, phagocytic mechanosensing of DAAM particles manifests as a binary choice between the rapid “gulping” of cargo and an alternative stalling response. Phagocytic cup maturation is typically associated with the clearance of F-actin from the center of the phagocyte-cargo interface (Cox, Tseng et al. 1999, Freeman, Goyette et al. 2016, Ostrowski, Freeman et al. 2019), which facilitates internalization of the newly formed phagosome into the cytoplasm. In videos of macrophages bound to soft DAAM particles, however, we typically observed a network of protrusive F-actin structures in the center of the contact for the entire duration of the experiment (40 minutes) (Supp Video 2, Fig 3J). This network was highly dynamic, with individual protrusions forming, moving, and dissolving on the timescale of minutes. The larger protrusions also physically distorted the cargo, creating micron-scale indentations on the particle surface (Supp Fig 4F). This complex and heterogeneous F-actin architecture contrasted sharply with the rapidly progressing phagocytic cups seen during stiff cargo engulfment, which were characterized by the strong accumulation of F-actin at the leading edge and concomitant F-actin clearance from the central domain (Supp Video 1).

To analyze these patterns in more detail, we mapped the F-actin signal of fixed, phalloidin-stained phagocytic cups onto the 3D rendered surfaces of their microparticle targets using a previously described analytical approach (Vorselen, Wang et al. 2020, Vorselen, Barger et al. 2021, de Jesus, Settle et al. 2023) (Supp Fig 4G). Our initial comparisons focused on early stage (< 60 % complete) interactions captured 5 minutes after BMDM-DAAM particle mixing. Contacts with stiff (18 kPa) and soft (0.6 kPa) particles both exhibited strong F-actin accumulation (Fig 3K), which we measured by dividing the mean F-actin intensity within the phagocytic cup by the mean intensity over the whole cell. The morphologies of these accumulations, however, were markedly different. Stiff particle contacts displayed a ring-like configuration, with strong F-actin enrichment at the front and clearance from base of the phagocytic cup, whereas soft particle contacts were more disorganized, with a poorly defined front and punctate structures at the base (Supp Fig 4G). To quantify these distinct patterns, we extracted two perpendicular intensity profiles through the center of the contact, then defined an actin clearance index as the ratio between mean F-actin intensity in the inner 50 % of the contact and mean intensity in the outer 50 % (Fig S3E). By this metric, early phagocytic cups that formed on stiff particles were significantly more “ring-like” (clearance ratio < 1) than their soft particle counterparts (Fig 3L). We applied the same analysis to soft particles cups that remained incomplete after 30 minutes, presumably due to phagocytic stalling. These contacts were even less ring-like than early stage soft particle interactions, suggesting that stalling results, at least in part, from the failure to clear central F-actin.

### *β*2 integrins respond differentially to stiff and soft cargo

We next sought to identify putative mediators of cargo mechanosensing. Reasoning that some of these molecules would be differentially phosphorylated in response to cargo rigidity, we performed phosphoproteomic analysis of BMDMs shortly after contact formation with IgG-coated DAAM-particles (Fig 4A). This experiment identified sets of peptides that were preferentially phosphorylated upon challenge with either soft (0.6 kPa) or stiff (18 kPa) targets (Supp Fig 5A-B). GO analysis of the phosphopeptides enhanced by engagement with stiff particles revealed associations with the F-actin cytoskeleton, membrane reorganization, as well as both Rac and Rho GTPase signaling (Supp Fig 5C). One of the most prominent of these peptides encompassed a.a. 757-766 of the *β*2 integrin subunit (also called CD18), containing phosphothreonine 760 (pT760) (Fig 4B-C, Supp Fig 5A). This finding interested us because Fc receptor engagement induces the affinity maturation of integrins (Jongstra-Bilen, Harrison et al. 2003, Chung, Serezani et al. 2008). In addition, the *β*2 integrin Mac-1, a heterodimer composed of CD18 and CD11b (*α*M), is the established phagocytic receptor for complement. *β*2 integrins have also been implicated in complement-independent forms of phagocytosis (Walbaum, Ambrosy et al. 2021, Vorselen, Kamber et al. 2022), although the basis for this involvement is poorly understood. Importantly, phosphorylation of human CD18 on T568, which is orthologous to mouse T760, has been associated with increased engagement of cytoskeletal adaptor molecules (Fagerholm, Hilden et al. 2005, Nurmi, Autero et al. 2007), hinting at a role for this protein in mechanosensing.

**Fig. 4.**
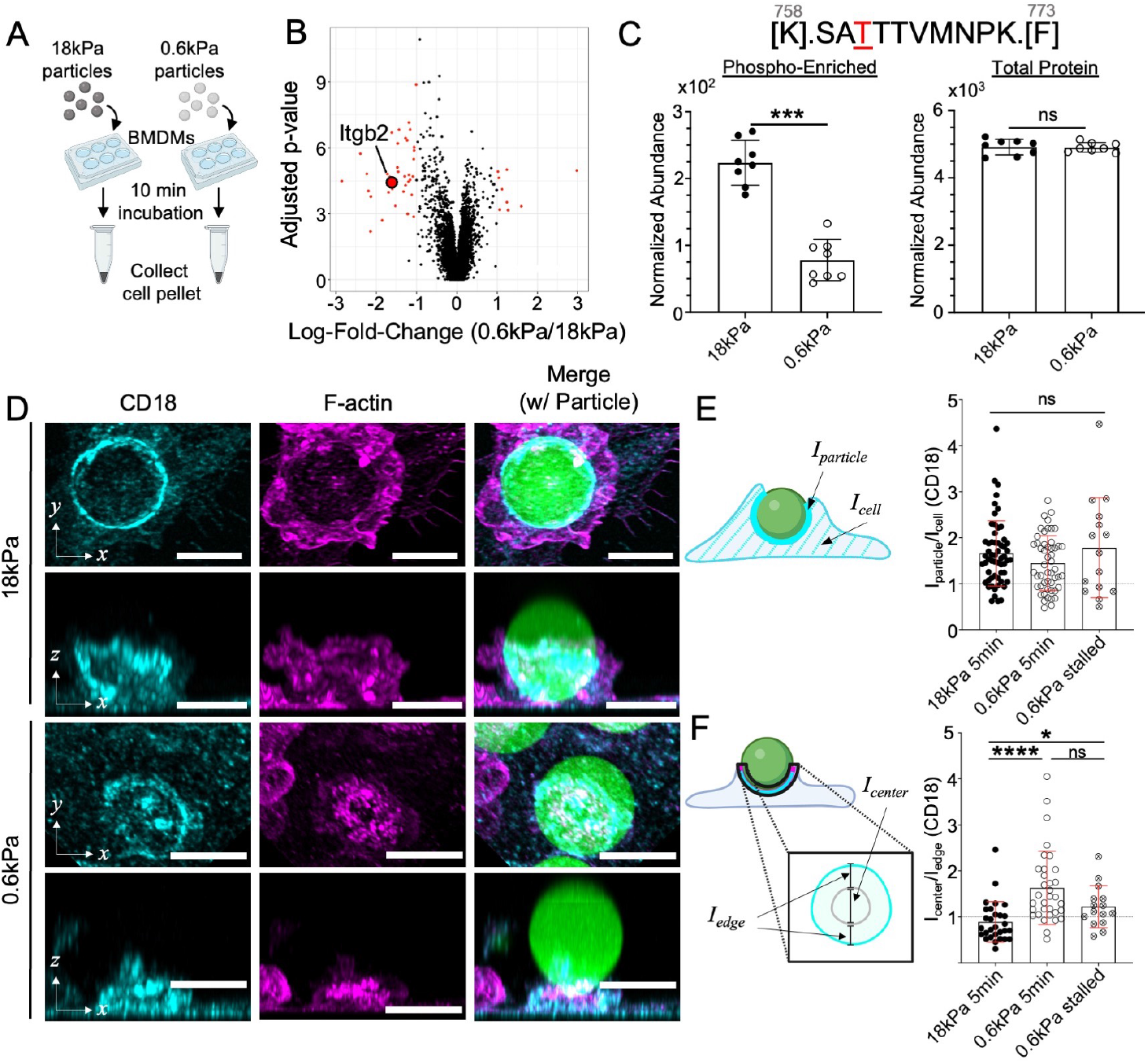
*β*2 integrins respond differently to stiff and soft targets. A-B) BMDMs were challenged with 18 kPa and 0.6 kPa IgG-coated DAAM particles and then subjected to phosphoproteomic analysis. A) Schematic diagram of the phosphoproteomic sample collection – BMDMs and microparticles were co-incubated for 10 minutes before transferring to ice, and cells were then lifted and pelleted at -80 °C. B) Volcano plot of the resulting phosphopeptides, with negative and positive log fold-change (LogFC) denoting enrichment with 18 kPa and 0.6 kPa particles, respectively. LogFC and significance were calculated using 8 independent replicates for each condition. C) Above: Sequence of Itgb2758-773, with phosphorylated threonine highlighted in red. Below left, normalized abundance values of Itgb2758-773 phosphopeptide. Below right, normalized abundance of all Itgb2 peptides without phospho-enrichment. n=8 replicates (independent macrophage differentiations from 3 pooled mice). Error bars denote SD. ^***^: adjusted p-value (Bonferroni) = 0.003. D-F) BMDMs were challenged with IgG-coated DAAM particles of different rigidities for various times, fixed and stained for F-actin and CD18, and then imaged by confocal microscopy. D) Representative images of DAAM particle-macrophage conjugates after 5 min incubation, stained with anti-CD18 antibody (Cyan) and phalloidin (F-actin, Magenta). Rows 1 and 3: top-down maximum intensity projections, Rows 2 and 4: side-view maximum intensity projections. Scale bars = 10 µm. E-F) CD18 accumulation (E) defined as the ratio of CD18 at the particle interface divided by total CD18 in the cell and CD18 clearance ratio (F) comparing CD18 signal at the periphery and the center of the phagocytic cup (see Materials and Methods) determined in phagocytic cups formed with 18 kPa (5 minutes) and 0.6 kPa (5 and 30 minutes) DAAM particles. In E and F, error bars denote SD. ^*^: p < 0.05, ^****^: p<0.0001, two-tailed Welch’s T-tests. n = 58, 53, and 15 cells for 18 kPa,0.6 kPa 5min, and 0.6 kPa 30min, respectively.

To investigate this possibility, we examined the localization of CD18 during phagocytosis. BMDMs were fixed 5 minutes after challenge with IgG-coated DAAM particles, and the resulting early-stage contacts were stained for F-actin and CD18 and then imaged (Fig. 4D). CD18 strongly accumulated in phagocytic cups engaging both stiff (18 kPa) and soft (600 Pa) particles, which was notable given that *β*2 integrins are not known to bind directly to IgG or FcR (Jongstra-Bilen, Harrison et al. 2003). To quantify this behavior, we calculated a CD18 accumulation ratio by dividing the mean fluorescence intensity in direct contact with the particle by the intensity across the whole cell. This analysis revealed marked and largely rigidity-independent CD18 recruitment to the phagocytic cup (Fig. 4E), consistent with a role for *β*2 integrins in the direct recognition and mechanical interrogation of cargo.

Next, we assessed the organization of CD18 at the BMDM-cargo interface using the clearance ratio methodology applied above to profile F-actin (Supp Fig 4G). Whereas CD18 formed ring-like structures on stiff particles (clearance ratio < 1), a more diffuse pattern prevailed during engagement with soft cargo, with appreciable accumulation at the center of the contact (Fig. 4F). Diffuse localization of CD18 was also observed in stalled phagocytic cups captured 30 minutes after mixing with soft particles (Fig. 4F). These data mirrored our results with F-actin, implying that CD18 and F-actin are recruited to at least some shared structures during phagocytosis. Consistent with this interpretation, we observed strong colocalization, determined by Pearson’s Correlation, between F-actin and CD18 in contacts formed with either stiff or soft cargo (Supp Fig 5D). This correspondence was not perfect, however, particularly in conjugates with stiff particles, where CD18 signal appeared to lag behind F-actin at the leading edge of the phagocytic cup. To quantify this behavior, we determined the distance between the F-actin and CD18 fronts in each contact, which we then normalized to the particle circumference. The resulting “Lag Distance” parameter was significantly larger in phagocytic cups containing 18 kPa particles (Supp Fig 5E), indicating that an F-actin rich leading edge lacking *β*2 integrins is a characteristic feature of stiff cargo engulfment. Hence, similar to our observations with F-actin, *β*2 integrin organization, but not accumulation, depends on cargo rigidity.

### *β*2 integrins couple cargo stiffness to the mechanism of uptake

To better understand the role of *β*2 integrins during phagocytosis, we turned to genetic loss-of-function approaches. First, we applied CRISPR/Cas9 technology to target exon 2 of the Itgb2 locus (encoding CD18) in HoxB8-ER myeloid progenitors, which we then differentiated into CD18 deficient (CD18-KO) macrophages (Fig 5A, Supp Fig 6A). Control macrophages were prepared in parallel using a nontargeting (NT) guide RNA. Upon challenge with IgG-coated DAAM particles, CD18-KO macrophages exhibited two phagocytosis defects. The first was a significant reduction in the uptake of both stiff (18 kPa) and soft (0.6 kPa) particles. Second, and more intriguing, was a complete lack of preference for stiff over soft targets, indicating that CD18-KO macrophages are incapable of cargo mechanosensing (Fig 5B). To examine this phenotype more closely, we derived and compared BMDMs from Itgb2-/- mice (Wilson, Ballantyne et al. 1993) and Itgb2+/littermate controls (Fig 5C). Itgb2-/BMDMs exhibited both a global decrease in phagocytic activity as well as reduced discrimination between stiff and soft targets (Fig 5D and Supp Fig 6B-C), similar to the behavior of CD18-KO HoxB8-ER macrophages. Importantly, the stiffness independent part of this phagocytosis defect was rescued by increasing the concentration of IgG used for DAAM particle coating to 100 pmol/M from 10 pmol/M (Fig 5E). High density IgG did not, however, restore mechanosensing; stiff and soft particles were still phagocytosed by Itgb2-/- BMDMs to the same extent (Fig 5E). These results indicate that *β*2 integrins both enable phagocytic mechanosensing and promote phagocytic throughput.

**Fig. 5.**
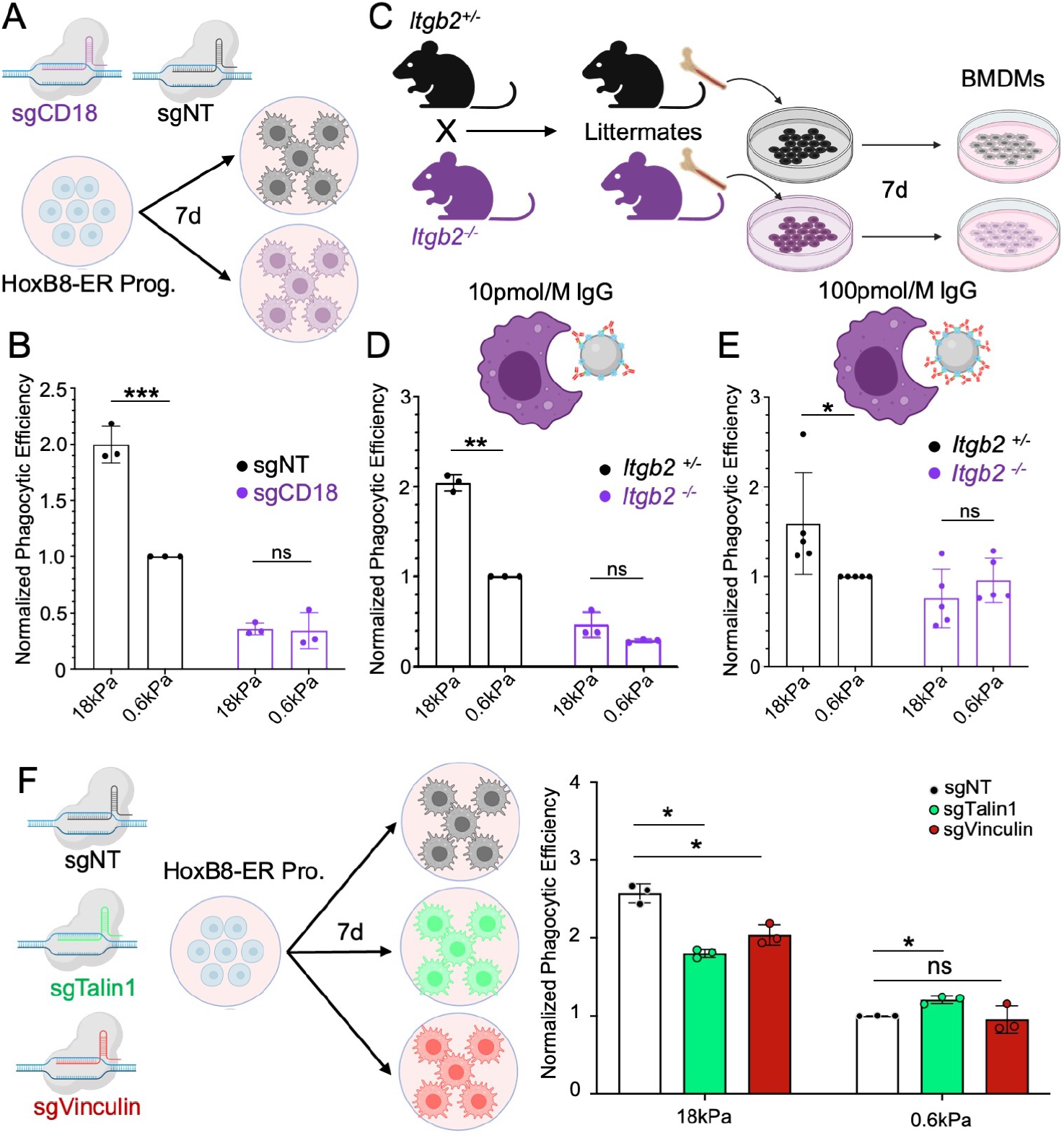
*β*2 integrin signaling is necessary for phagocytic mechanosensing. A) HoxB8-ER conditionally immortalized progenitors were transduced with either a non-targeting sgRNA or sgRNA targeting exon 2 of CD18, and then differentiated into macrophages. B) sgCD18 HoxB8-ER macrophages and sgNT controls were challenged with IgG-coated 18 kPa and 0.6 kPa DAAM particles. Graph shows normalized phagocytic efficiency, defined as the ratio of % phagocytic cells relative to the sgNT, 0.6 kPa sample. n = 3 biological replicates from independent differentiations. ^***^: p<0.001, ns: not significant, two-tailed Ratio Paired t-test. C) Diagram schematizing the preparation of BMDMs from Itgb2-/- mice. Heterozygous (Itgb2+/-) and homozygous knockout mice (Itgb2-/-) mice were bred and littermate F1 offspring of each genotype were compared in each biological replicate. D-E) Itgb2-/- BMDMs and Itgb2+/- controls were challenged with 18 kPa and 0.6 kPa DAAM particles coated with 10 pmol/M (D) and 100 pmol/M (E) IgG. Graphs show phagocytic efficiency normalized against the Itgb2+/-, 0.6 kPa sample. n = 3 biological replicates for each genotype. ^**^: p<0.01, ^*^: p<0.05, ns: not significant, two-tailed Ratio Paired t-test. F) HoxB8-ER conditionally immortalized progenitors were transduced with either a non-targeting sgRNA or sgRNA targeting Talin1 or Vinculin, and then differentiated into macrophages. These cells were then challenged with IgG-coated 18 kPa and 0.6 kPa DAAM particles. Graphs show phagocytic efficiency normalized against the sgNT, 0.6 kPa sample. n = 3 biological replicates for each genotype. ^*^: p<0.05, ns: not significant, two-tailed Paired Ratio T-tests. All error bars denote SD.

Integrins mediate mechanosensing through the scaffolding proteins Talin and Vinculin, which form mechanically driven complexes that couple integrins to the F-actin cytoskeleton and initiate downstream signaling (Kim, Ye et al. 2011, Bays and DeMali 2017). To evaluate the importance of Talin and Vinculin for cargo mechanosensing, we prepared HoxB8-ER macrophages lacking each protein using the CRISPR/Cas9 targeting approach described above (Fig 5F, Supp Fig 6D). Depletion of either molecule significantly reduced the phagocytosis of stiff (18 kPa) DAAM particles without affecting soft (0.6 kPa) particle uptake (Fig 5F). Taken together with the CD18-focused experiments described above, these data strongly suggest that the formation of *β*2 integrin adhesions drives the accelerated consumption of stiff cargo.

We reasoned that *β*2 integrin adhesions might trigger “gulping” phagocytosis of stiff particles by inducing the annular F-actin architecture characteristic of this uptake mechanism. To test this hypothesis, we challenged Itgb2-/- BMDMs and Itgb2+/- controls with IgG-coated DAAM particles of differing stiffness, then assessed phagocytic cup progress after 5 minutes by fixing and staining for F-actin (Fig 6A-D). High concentration IgG coating was used in this experiment to increase conjugate formation in the Itgb2-/- samples. As expected, Itgb2+/- BMDMs exhibited strong, predominantly ring-like F-actin enrichment in phagocytic cups containing stiff (18 kPa) particles (Fig 6A, E-F). This characteristic accumulation pattern was almost completely abrogated in Itgb2-/- BMDMs (Fig 6C, E-F), indicating both that *β*2 integrins control F-actin configuration on stiff cargo and that this configuration is essential for “gulping” phagocytosis.

**Fig. 6.**
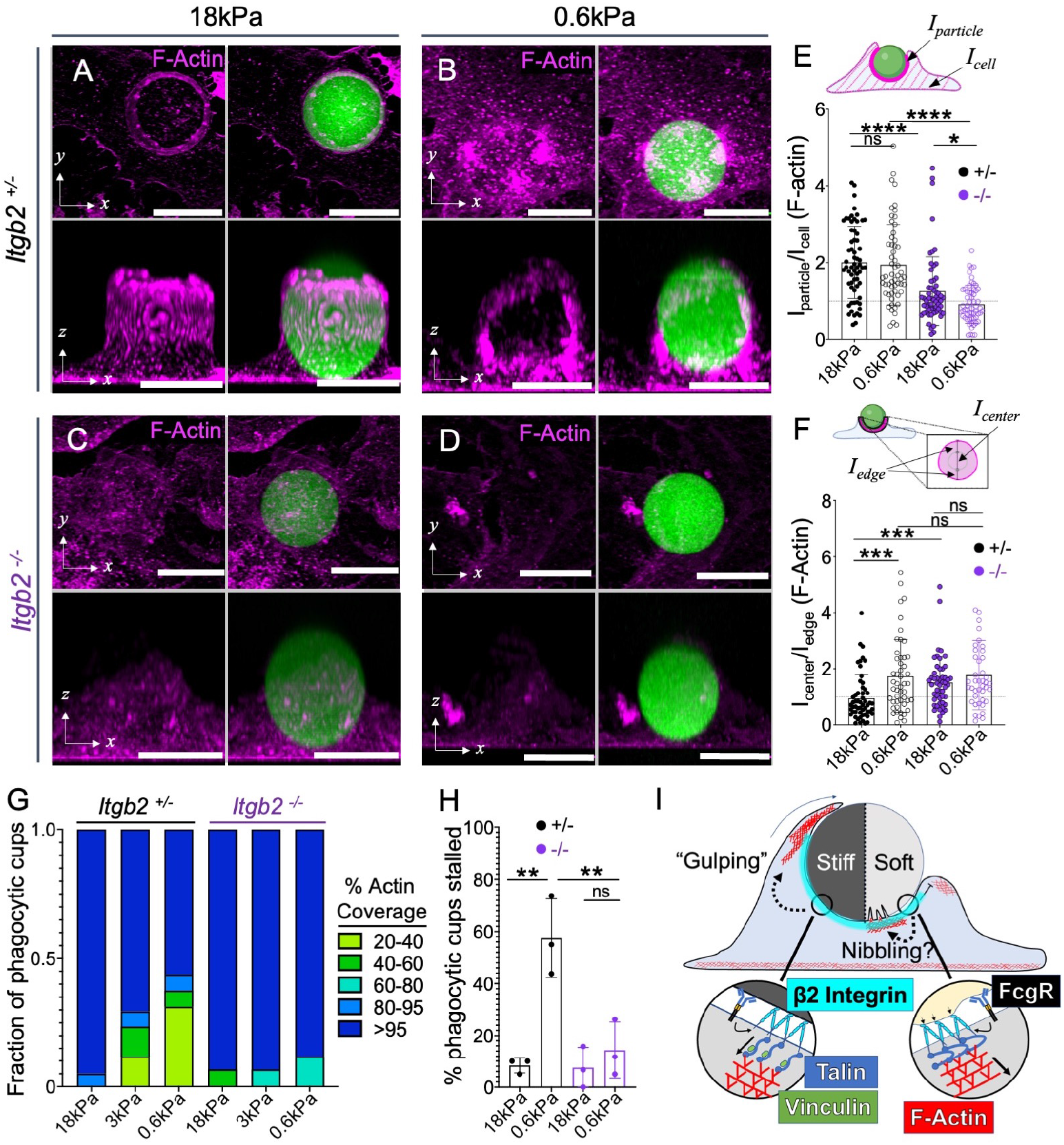
*β*2 integrins drive distinct responses to stiff and soft cargo. Itgb2-/- BMDMs and Itgb2+/- controls were challenged for various times with DAAM particles of differing rigidity coated with 100 pmol/M IgG, fixed and stained for F-actin, and imaged by confocal microscopy. A-D) Representative maximum z-projections (top) and y-projections (bottom) of Itgb2+/- (A and B) and Itgb2-/- (C and D) BMDMs in the act of engulfing 18 kPa (A and C) and 0.6 kPa (B and D) DAAM particles (green). F-actin is visualized in magenta. Scale bars = 10 µm. E) F-actin accumulation (see Materials and Methods) for phagocytic cups graphed a function of particle stiffness and Itgb2 genotype. n57 events. ^****^: p<0.0001, ^*^:, p<0.05, two-tailed Welch’s t-test. F) F-actin clearance ratio (see Materials and Methods) for phagocytic cups graphed as a function of particle stiffness and Itgb2 genotype. ^***^: p<0.001, two-tailed Welch’s t-test. G) F-actin coverage after 30 min incubation, determined for Itgb2-/- BMDMs and Itgb2+/- controls. Values are sorted into five bins, with colors denoting the fraction of events falling within each bin. Representative bars from one biological replicate are shown, from 20 events per condition. H) Fraction of events that are 20-95 % complete after 30 min co-incubation of DAAM particles with BMDMs. n = 3 biological replicates (litter-mate pairs, 20 events per condition per replicate). ^**^: p<0.01, Paired t-test. All error bars denote SD. I) *β*2 integrins mediate a mechanosensitive checkpoint that initiates actin accumulation at the phagocytic cup, thereby enabling the rapid “gulping” of stiff cargo (left) and the formation of central actin protrusions to engage soft cargo, potentially as part of a “nibbling” response (right).

Interestingly, Itgb2 deficiency disrupted F-actin recruitment to soft (600 Pa) particles, as well. Whereas Itgb2+/- BMDMs formed F-actin rich phagocytic cups on this cargo without a central clearance (Fig 6B, E-F), the corresponding Itgb2-/- interactions lacked any discernable F-actin enrichment at all (Fig 6D-F). These observations raised the possibility that the phagocytic stalling induced by soft cargo does not simply represent a failure to gulp, but rather a second, distinct type of *β*2 integrin-dependent uptake response. To investigate this idea, we compared phagocytic cup progression in Itgb2-/- and Itgb2+/- BMDMs 30 minutes after challenge with DAAM particles of varying stiffness. Stalling by Itgb2+/- BMDMs increased with decreasing cargo rigidity (Fig 6G-H), mirroring the results we obtained using particles coated with low density IgG (see Fig 3H-I). In striking contrast, Itgb2-/- BMDMs exhibited little to no stalling in any stiffness regime (Fig. 6G-H). With even the softest (0.6 kPa) microparticles, nearly all conjugates achieved complete engulfment by 30 minutes. Together, these unexpected results indicate that *β*2 integrins play instructive roles in both the gulping phagocytosis of stiff cargo and the stalling response to soft targets.

## Discussion

In the present study, we applied novel DAAM particle technology to study phagocytic mechanotransduction in macrophages independently of confounding factors such as cargo size and ligand presentation. Using this approach, we found that multiple macrophage subtypes respond to cargo rigidity and that they do so in the context of both IgG- and PtdS-induced uptake. For IgG-dependent phagocytosis, cargo mechanosensing manifested as a choice between rapid, “gulping” uptake, which was frequently observed on stiff DAAM particles, and phagocytic stalling, which was more prevalent with soft particles. These two forms of engagement differed not only in their dynamics but also in their underlying F-actin architectures: gulping interfaces were characterized by a strong peripheral ring of F-actin, whereas stalled interfaces exhibited F-actin accumulation both in the periphery and in punctate structures closer to the center of the contact. Similar stiffness-induced effects on cup morphology were observed in a previous study of “frustrated” phagocytosis on planar substrates (Hu, Li et al. 2023). Importantly, we found that gulping and stalling responses both required *β*2 integrins. In the absence of CD18, each form of uptake was replaced by a third mode in which little to no F-actin accumulated at the phagocytic cup. Based on these results, we conclude that macrophages apply a *β*2 integrin-dependent mechanical checkpoint to tailor phagocytic mechanism to the deformability of their targets (Fig 6I). That CD18 deletion reduces stalling on soft DAAM particles strongly suggests that stalling is an actively induced process and not just a failed attempt at gulping. This raises the question of what stalling actually represents and how this mechanism would manifest in interactions with more physiologically relevant cargos. The central F-actin protrusions that form in interactions between macrophages and soft DAAM particles are reminiscent of the projections that mediate phagocytic “nibbling”, or trogocytosis (Velmurugan, Challa et al. 2016, Vorselen, Kamber et al. 2022, Zhao, Zhang et al. 2022). This piecemeal engulfment process is thought to be important for the uptake of apoptotic corpses and tumor cells, as well as axonal pruning by microglia (Brouckaert, Kalai et al. 2004, Matlung, Babes et al. 2018, Weinhard, di Bartolomei et al. 2018, Lim and Ruthazer 2021, Dooling, Andrechak et al. 2023). In each of these cases, the phagocytic target is orders of magnitude softer than typical bacterial and fungal cargos. In light of our results, it is tempting to speculate that deformability serves as a physical trigger to induce phagocytic nibbling, which would facilitate the consumption of physiological targets like apoptotic cells. When faced with an equally deformable but unfragmentable DAAM particle, however, attempted nibbling would instead lead to the unproductive, phagocytic stalling that we observed.

In the absence of *β*2 integrins, we found that macrophages employed a third form of phagocytosis characterized by the absence of both F-actin enrichment and mechanosensing. This rudimentary mode of uptake was particularly sensitive to ligand density, implying that abundant phagocytic receptor engagement can drive engulfment in the absence of integrin-induced adhesion and cytoskeletal remodeling. These results are intriguing in light of recent work showing that that Itgb2-/- macrophages form aberrantly elongated phagocytic cups with IgG-coated red blood cells that are nevertheless capable of completing phagocytosis (Walbaum, Ambrosy et al. 2021). They are also consistent with prior work indicating that integrin binding is critical for cup formation when phagocytic ligands are sparsely clustered (Freeman, Goyette et al. 2016). Although further studies will be required to understand the mechanistic basis of this rudimentary uptake pathway, its existence strongly suggests that *β*2 integrins do not enable phagocytosis per se, but rather the application of distinct phagocytotic strategies that best match the specific mechanical features of the cargo.

Previous studies have documented integrin dependent mechanosensing in the context of focal adhesions and podosomes (Collin, Na et al. 2008, Friedland, Lee et al. 2009, Case and Waterman 2015, Driscoll, Bidone et al. 2021), both of which are F-actin rich assemblies that impart force against extracellular entities. Within these structures, the stress/strain relationship between force exertion and environmental deformation is interpreted via outside-in integrin signaling, ultimately resulting in stiffness dependent cellular responses. It is tempting to speculate that cargo mechanosensing by macrophages proceeds by an analogous mechanism in which *β*2 integrin mechanotransduction couples the degree of target deformation to the appropriate mode of uptake. Thus, the inability to deform cargo would induce phagocytic gulping, whereas strong distortion would prime the alternative uptake mechanism discussed above. The F-actin rich protrusions that we and others (Ostrowski, Freeman et al. 2019, Tertrais, Bigot et al. 2021, Vorselen, Barger et al. 2021, Vorselen, Kamber et al. 2022) have observed in nascent phagocytic cups seem likely candidates for this function because they readily deform target surfaces. From a design perspective, mechanosensing structures would not need to be distinct from the protrusions used to fragment and/or engulf cargo. Indeed, investing the same protrusion with both uptake and mechanosensory potential would better equip a macrophage to change its phagocytic strategy in response to a particularly dynamic or heterogeneous meal.

*β*2 integrins are not known to interact directly with IgG and PtdS, raising the question of what these receptors recognize on the IgG- and PtdS-coated DAAM particles used in this study. It is formally possible that *β*2 integrins influence the mechanosensing of this cargo indirectly by establishing strong contact with underlying substrate ECM. The fact that CD18 strongly localizes to the phagocytic cup, however, regardless of cargo stiffness, argues against this hypothesis. Certain integrins, including Mac-1 and X*β*2, are notably promiscuous, exhibiting appreciable affinity for denatured proteins and even uncoated plastic (Yakubenko, Lishko et al. 2002, Lamers, Plüss et al. 2021). As such, it seems most likely that *β*2 integrins bind directly either to the polyacrylamide matrix of the DAAM particle itself or, alternatively, to serum derived proteins adsorbed onto the particle surface. In physiological settings, we expect that the promiscuity of *β*2 integrins, combined with the molecular heterogeneity of biological targets, would enable some form of direct integrin recognition in most phagocyte-cargo pairings.

The specific CD18 phosphorylation event identified by our proteomic analysis (pT760 in mice, pT758 in humans) occurs in response to TCR ligands, phorbol ester, and chemokines (Fagerholm, Varis et al. 2006, Jahan, Madhavan et al. 2018). This creates a recognition site for 14-3-3 proteins, which bind to the CD18 tail and thereby alter the composition of the adhesion complex. The functional consequences this process remain somewhat unclear; while some studies indicate that 14-3-3 binding promotes cell adhesion through the activation of Rho family GTPases, it has also been shown to disrupt interactions between CD18 and Talin (Fagerholm, Hilden et al. 2005, Nurmi, Autero et al. 2007, Takala, Nurminen et al. 2008), which could attenuate integrin function. Here, we demonstrate that T760 phosphorylation is enhanced by target rigidity. This mechanoresponsiveness may simply reflect stronger engagement of *β*2 integrins, which would be expected to boost outside-in integrin signaling as well as any downstream feedback inhibition. It is also possible, however, that T760 phosphorylation plays a specific role in promoting the phagocytic gulping of stiff cargo. Targeted mechanistic studies will be required to resolve this issue.

In recent years, the potential of phagocyte-based immunotherapy has motivated intensive investigation of its governing mechanisms. Much attention has focused on the “eat” versus “don’t eat” choice point, no doubt reflecting the success of strategies targeting “don’t eat me” signals like CD47 (Maute, Xu et al. 2022). This critical decision, however, is just the first of many on the path to cargo uptake. Once the macrophage commits to phagocytosis, the key question becomes how to consume the cargo. Our results clearly demonstrate that this next stage is subject to complex biomechanical regulation. Decoding the molecular logic that controls this process could unlock new strategies for modulating phagocytic activity in translational settings.

## Data Availability

Raw proteomic LC/MS data and imaging files are available upon request from the corresponding author (Morgan Huse).

## Code Availability

All custom MATLAB scripts are available upon request from the corresponding author (Morgan Huse).

## ACKNOWLEDGEMENTS

We thank the staff of the Research Animal Resource Center, The Flow Cytometry and Cell Sorting Core, Molecular Cytology Core, and the Proteomics Core at MSKCC for their assistance. We also thank the members of the Perry and Huse labs for advice and technical assistance. We gratefully acknowledge David B. Sykes for the generous gift of HoxB8-ER plasmids and cell lines. Supported in part by the NIH (R01-AI087644 to M.H.), the MSKCC Basic Research Innovation Award Program (M. H.), the National Science Foundation Graduate Research Fellowship program (A.H.S.), and the Center for Experimental Immuno-Oncology at MSKCC (A.H.S.).

## AUTHOR CONTRIBUTIONS

A.H.S, D.V., J.S.A.P, and M.H. conceived of the project. A.H.S and M.H. analyzed data, generated all figures, and wrote the manuscript. A.H.S., B.Y.W, M.M.D.J., L.S., Z.W., Z.L., M.M.M., performed experiments and/or analysis for the manuscript. B.Y.W., M.M.D.J., E.C., Y.R., R.C.H., D.V. and J.S.A.P provided key expertise in experimental design and interpretation.

## Supplementary Material

**Supplementary Video 1**: Live imaging of stiff cargo phagocytosis.

Representative time-lapse microscopy videos at 63× magnification of HoxB8-ER derived macrophages transduced with f-tractin mCherry (magenta) phagocytosing 18 kPa IgG-coated DAAM particles (green, yellow arrows). All three examples display rapid uptake in less than 5 min. For each example, a 3D maximum intensity projection (rendered in Imaris) is displayed followed by orthogonal angles, slices chosen to capture as much of the phagocytic process as possible.

**Supplementary Video 2: Live imaging of phagocytic stalling on soft cargo**. Representative time-lapse microscopy videos at 63× magnification of HoxB8-ER derived macrophages transduced with f-tractin mCherry (magenta) interacting with 0.6 kPa IgG-coated DAAM particles (green, yellow arrows). In both examples, phagocytosis does not complete.

**Supplementary Video 3: Live imaging of soft cargo phagocytosis**. An additional video showing a HoxB8-ER macrophage expressing f-tractin mCherry (magenta) interacting with a 0.6kPa IgG-coated DAAM particle (green). At the start of the time-lapse, engulfment had proceeded to 75 % completion, but remained incomplete for an additional 8 min while exhibiting stereotypical features of stalled phagocytosis (central actin protrusions, particle deformation). Shortly thereafter, however, phagocytosis completed. An oblique slice of the f-tractin-mCherry signal along the axis of the phagocytic cup is shown for clarity.

**Fig. S1.**
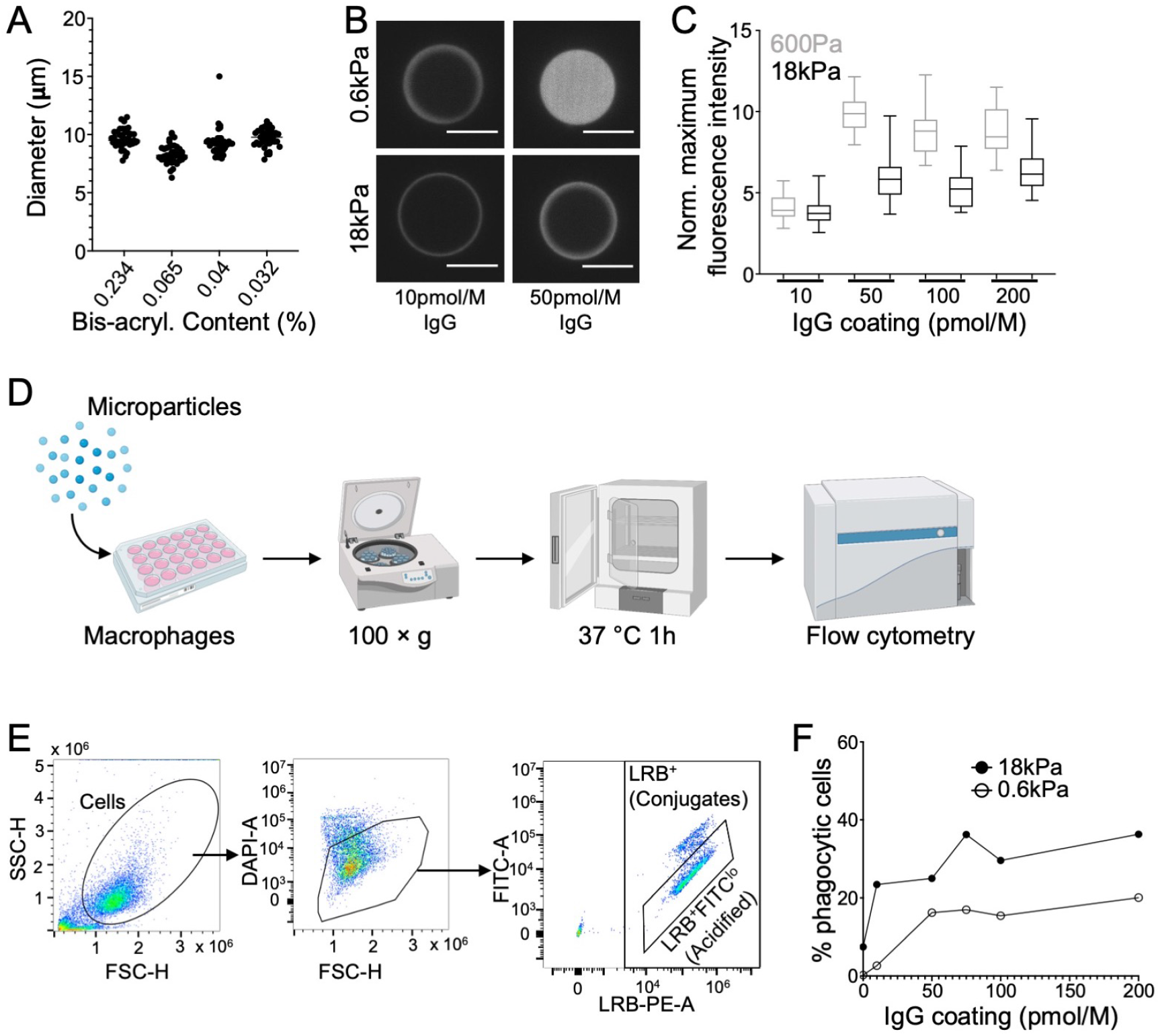
A hydrogel microparticle system to interrogate phagocytic mechanosensing. A) Microparticle diameter measured by confocal microscopy. n = 39 particles per condition. B) Confocal microscopy images of IgG-coated particles counter-stained with anti-Mouse IgG. Scale bars = 10 µm. C) Quantification of effective IgG coating as a function of coating concentration. Maximum fluorescence intensity is normalized in each image to an AccuCheck normalization bead within the same field of view (not shown). D) Schematic diagram for flow cytometric phagocytosis assay workflow. E) Representative gates for flow cytometric phagocytosis assay. Phagocytic efficiency is measured as LRB+FITClo events / all live cells. F) Phagocytic efficiency of stiff (18 kPa) and soft (0.6 kPa) DAAM particles, graphed as a function of IgG coating concentration.

**Fig. S2.**
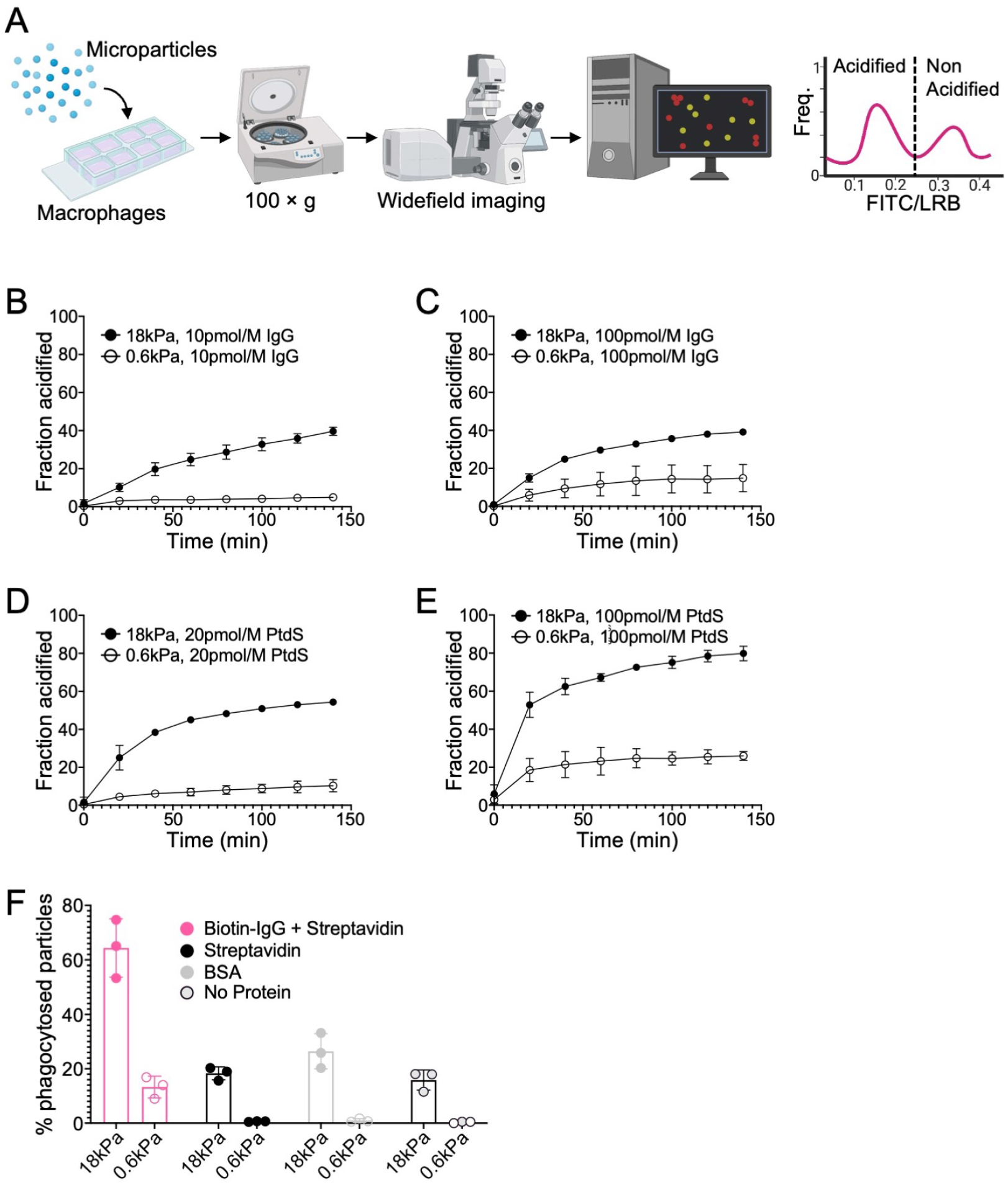
Cargo rigidity promotes phagocytic uptake. A) Schematic diagram for wide-field microscopy-based phagocytosis assay and quantification. An automated analysis script is used to identify particles, measure FITC/LRB ratios, and categorize particles as acidified or non-acidified. B-E) BMDMs were challenged with 18 kPa or 0.6 kPa DAAM particles coated with IgG or PtdS, and phagocytic efficiency was measured over time by widefield live microscopy. n=3 technical replicates from values averaged across four fields of view per replicate. F) BMDMs were imaged together with 18 kPa or soft 0.6 kPa DAAM particles that had been coated as follows. Biotin-IgG + Streptavidin: standard coating protocol with 250 pmol/M Streptavidin and 10pmol/M IgG + 50 µM FITC/LRB. Streptavidin: Particles conjugated with 250 pmol/M Streptavidin + 50 µM FITC/LRB. BSA: Particles conjugated with 250 pmol/M BSA + 50 µM FITC/LRB, No Protein: Particles conjugated with 50 µM FITC/LRB. Phagocytosis was quantified as the fraction of particles acidified per field of view after 2 h. Each point represents the mean of four fields of view in one technical replicate. Error bars denote SD, determined from 3 technical replicates.

**Fig. S3.**
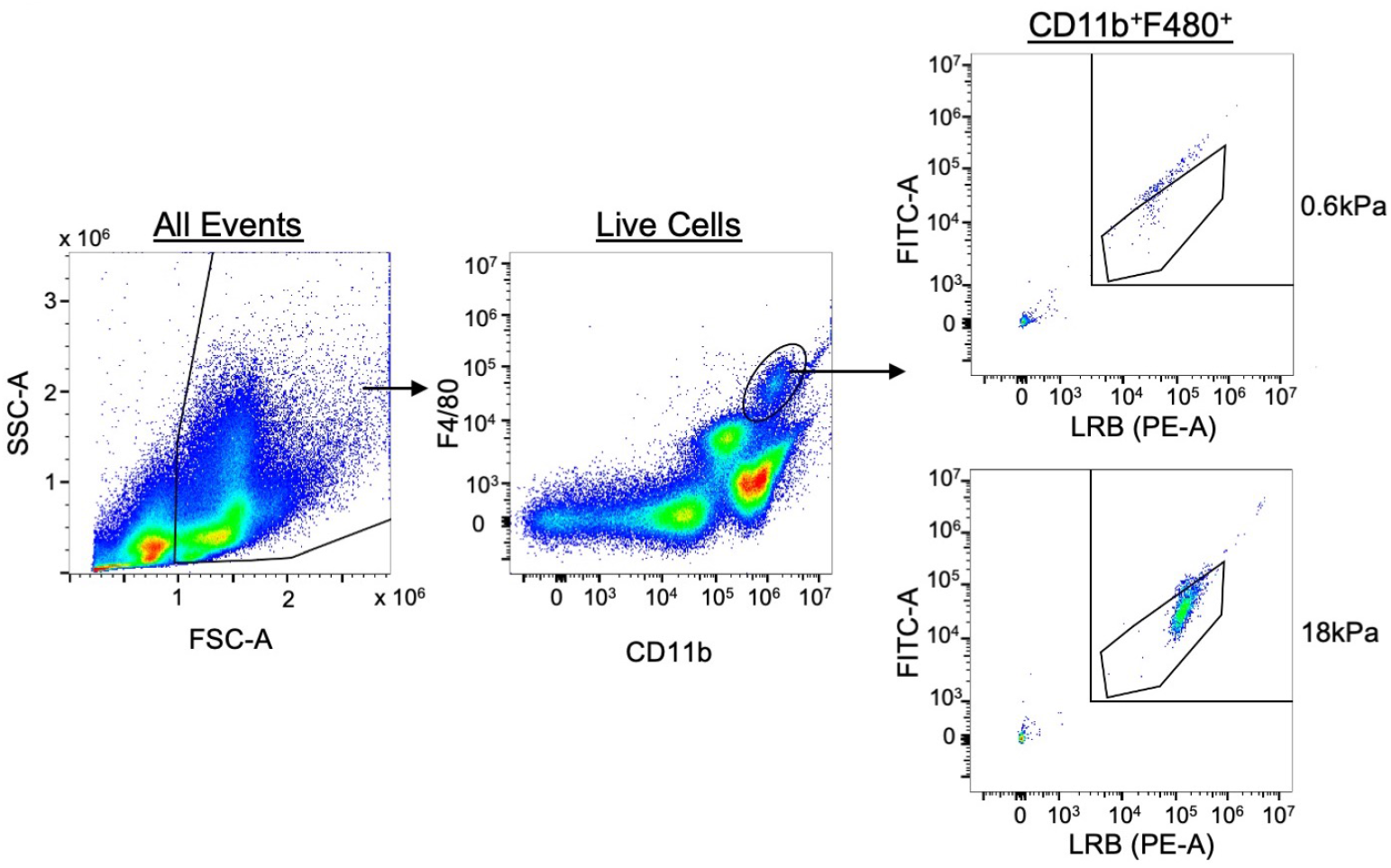
Gating strategy for in vivo peritoneal macrophage phagocytosis assay. Macrophages were identified as CD11b+F4/80+ and phagocytic cells as LRB+FITClo.

**Fig. S4.**
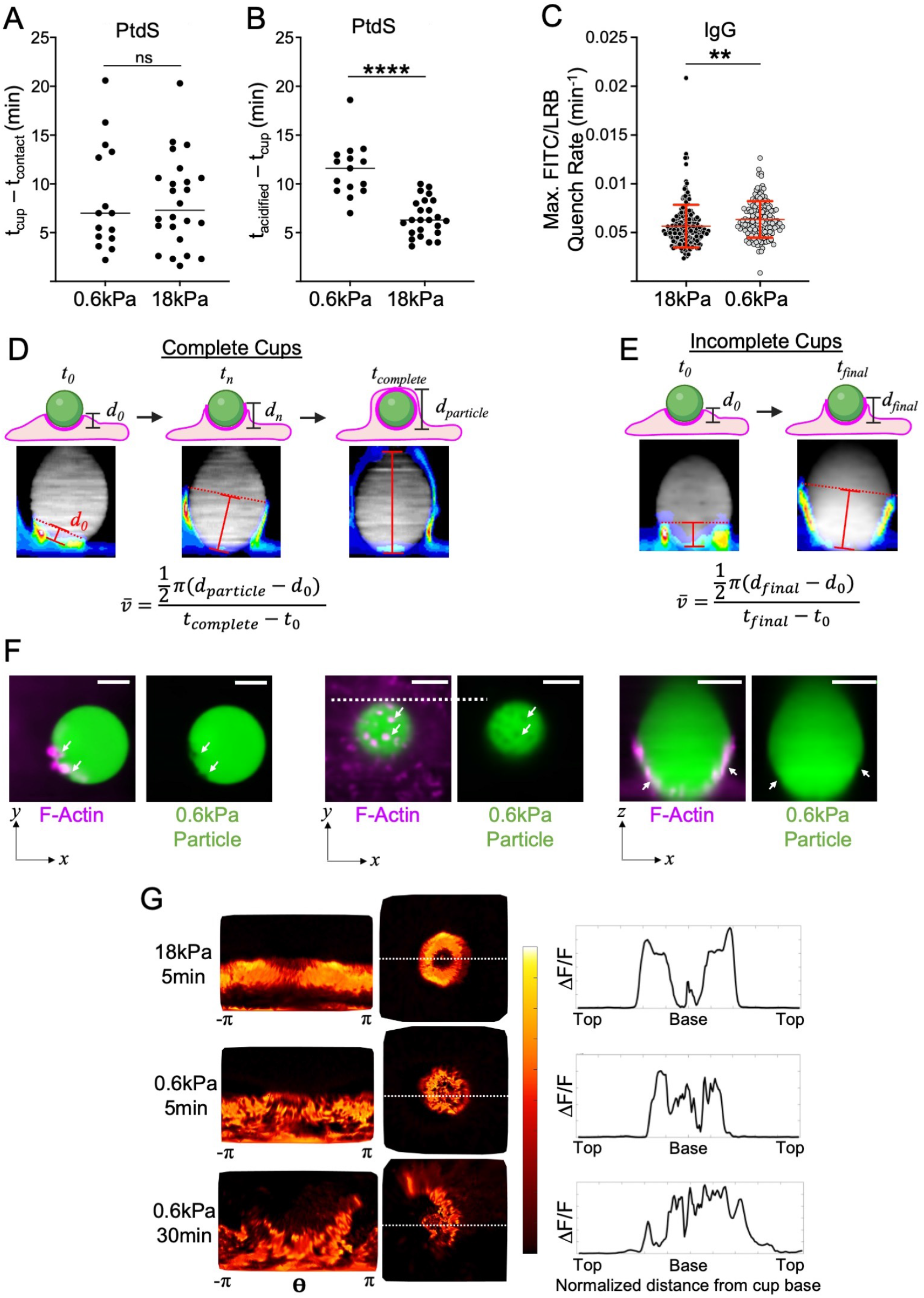
Phagocytosis of soft targets is slow and prone to stalling. A-B) BMDMs were challenged with 18 kPa or 0.6 kPa PtdS-coated DAAM particles and imaged by widefield videomicroscopy. A) Recognition time, defined as time difference between cup formation and first contact, n15 cells, ns: not significant. Two-tailed Welch’s t-test. B) Total acidification time, defined as the difference between particle acidification and cup formation. n=25 cells each condition, ^****^: p<0.0001, two-tailed Welch’s T-test. In A and B, mean values are indicated by horizontal lines in each column. C) BMDMs were challenged with 18 kPa or 0.6 kPa IgG-coated DAAM particles and imaged by widefield videomicroscopy. Graph shows maximum acidification rate, measured as the maximum rate of decrease in FITC/LRB ratio in each particle/cell interaction. n 147 cells, pooled from 3 technical replicates. Error bars indicate SD. ^**^: p < 0.01, two-tailed Welch’s T-test. D-F) HoxB8-ER macrophages expressing f-tractin-mCherry were challenged with IgG-coated DAAM particles and imaged live by spinning disk confocal microscopy. D-E) Schematic diagram for quantification of cup advancement in live movies (see Materials and Methods), showing the protocol for cups that complete (D) and for cups that stall (E). F) Representative images of 0.6 kPa contacts highlighting interfacial F-actin protrusion. Left, z-slice through center of a particle with protrusion-induced distortions indicated by white arrows. Center, z-slice near the base of a particle, with upward protrusions and particle indentations indicated by white arrows. Right, Side-view of a particle with indentations in the particle base indicated by white arrows. Scale bars = 5µm. G) BMDMs were challenged with IgG-coated DAAM particles of different rigidities for various times, fixed and stained for F-actin, and then imaged by confocal microscopy. Center, stereographic views of representative phagocytic cups, formed on particles of the indicated stiffness at the indicated time. Views are centered on the base of the cup and colored by local actin intensity. Left, world map views of each cup, aligned such that the base of the phagocytic cup is downward. Right, linescan profiles of normalized F-actin across each cup, determined using the dotted lines in the central images.

**Fig. S5.**
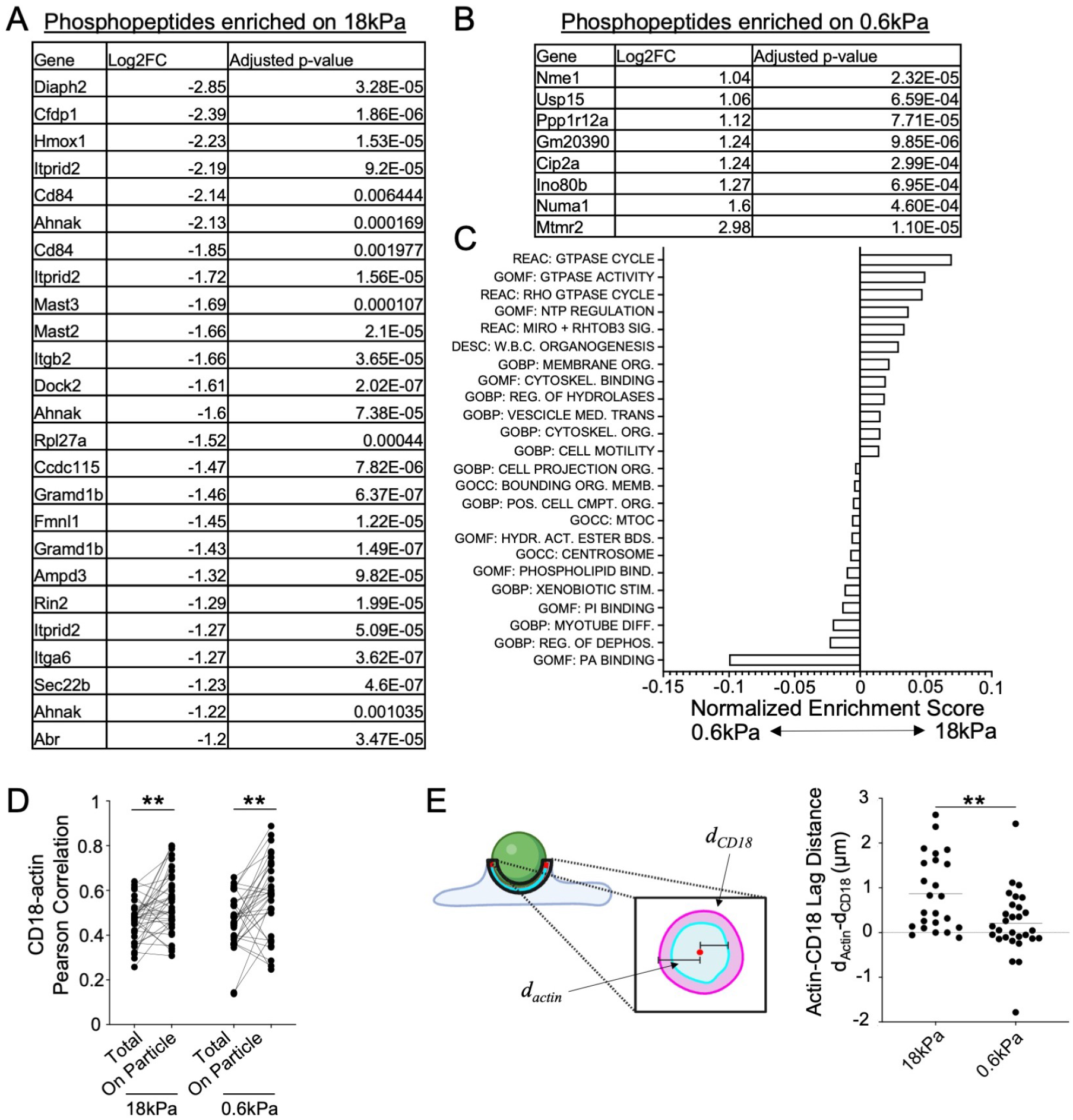
*β*2 integrins respond differently to stiff and soft targets. A-C) BMDMs were challenged with stiff (18 kPa) and soft (0.6 kPa) IgG-coated DAAM particles and then subjected to phosphoproteomic analysis. Tables show genes with phosphopeptides enriched in 18 kPa sample with Log2FC < -1.2 (A) or in the 0.6 kPa sample with Log2FC > 1 (B). Genes with multiple rows contain multiple enriched peptides. See Materials and Methods for statistical analysis. C) Pathway enrichment analysis of the phosphopeptide data, showing all significantly enriched gene groups. Databases queried: REAC: Reactome Pathway Database, GOMF: Gene Ontology Molecular Function, GOBP: Gene Ontology Biological Process, GOCC: Gene Ontology Cellular Compartment, DESC: DESCARTES. D-E) BMDMs were challenged with IgG-coated DAAM particles of different rigidities for various times, fixed and stained for F-actin and CD18, and then imaged by confocal microscopy. D) Colocalization between F-actin and CD18 (measured by Pearson’s Correlation Coefficient) in the whole cell body (Total) and in the phagocytic cup (On Particle). ^**^: p<0.01, two-tailed Paired t-test. E) CD18-actin lag distance, measuring the distance between F-actin and CD18 signal at the leading edge of the phagocytic cup (see Materials and Methods), determined on 18kPa and 0.6kPa particles. Mean values are indicated by the horizontal line in each column. ^**^: p<0.01, two-tailed Welch’s t-test.

**Fig. S6.**
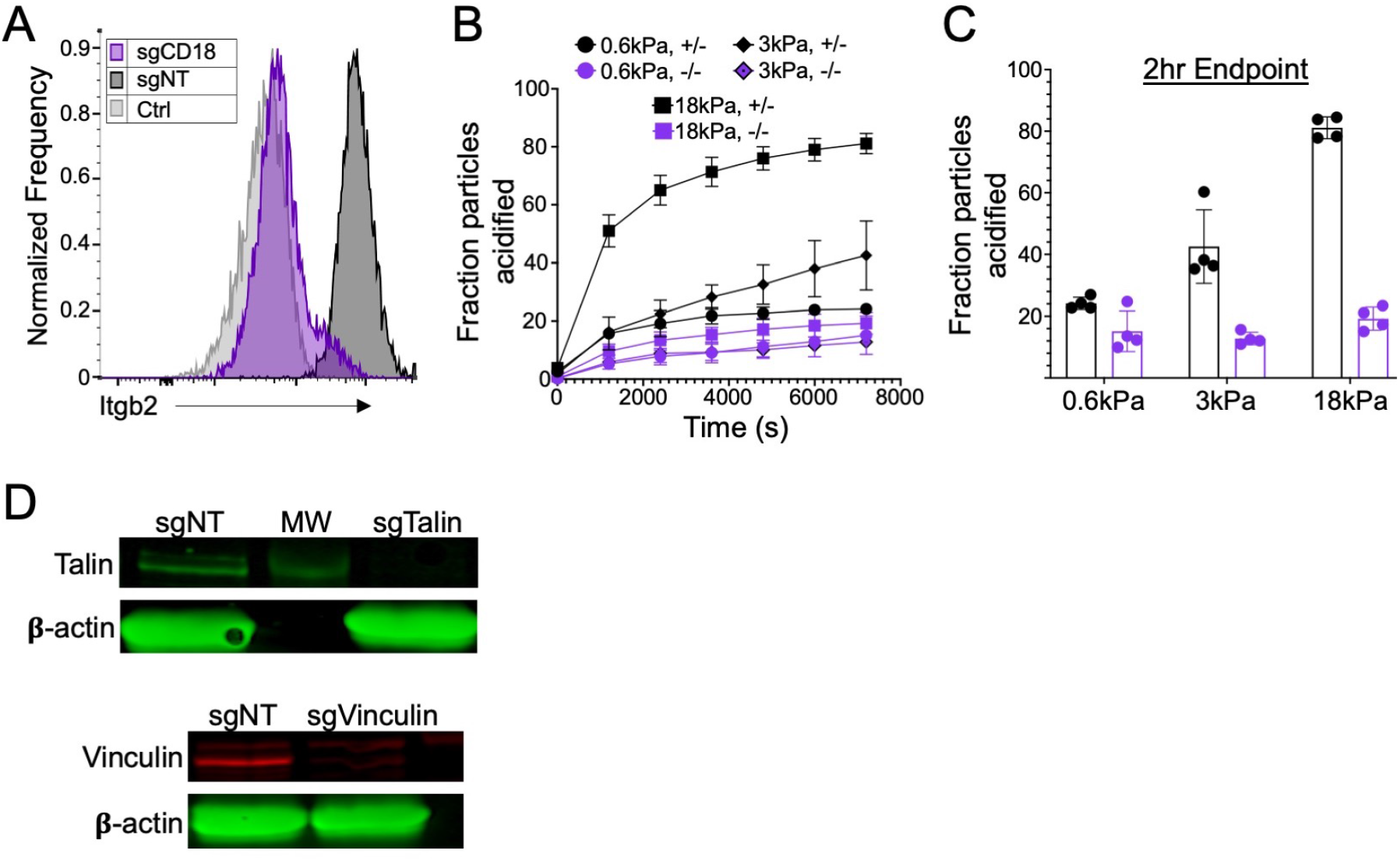
*β*2 integrin signaling is necessary for phagocytic mechanosensing. A) Representative histogram from flow cytometric analysis of CD18 expression in HoxB8-ER derived macrophages transduced with either sgCD18 (purple) or sgNT (black) guide RNAs, compared to a non-stained control (Ctrl). B-C) Itgb2-/- BMDMs and Itgb2+/- controls were imaged together with DAAM particles of differing rigidity coated with 10 pmol/M IgG. B) Kinetic analysis of phagocytosis, using automated image analysis to identify acidified particles (see Materials and Methods). n = 3 technical replicates, each from 4 analyzed fields of view. C) Phagocytosis was quantified as the fraction of particles acidified per field of view after 2 h. n = 4 technical replicates, each from 4 analyzed fields of view. Error bars in B and C indicate SD. D) Western blot analysis of HoxB8-ER derived macrophages transduced with sgNT, sgTalin, and sgVinculin. Talin expression is shown above and Vinculin expression below. *β*-actin served as a loading control. MW= Molecular weight 250k band. See Materials and Methods for antibodies used.

## Materials and Methods

### Plasmids and cell lines

For lentiviral transduction of f-tractin mCherry (plVX-f-Tractin-mCherry), the full f-tractin-mCherry coding sequence (from C1-F-tractin-mCherry, Addgene #155218) was subcloned into the lentiviral expression vector pLVX-M-puro (Takara 632164). For CRISPR-KO experiments, oligos for sequences targeting CD18 (CCTGTTCTAAGTCAGCGCCC), Talin (GCTTGGCTTGTGAGGCCAGT), Vinculin (GCAC-CTGGTGATTATGCACG), and non-targeting control (GCGAGGTATTCGGCTCCGCG) were annealed and cloned into lenti-sgRNA Blast (Addgene #104993). E0771 cells lines stably expressing a doxycycline-inducible MRTF-A construct were generated as previously described by sequential transduction with rTTA, TGL and either an MRTF-A or empty vector control expression construct(Tello-Lafoz, Srpan et al. 2021). B16-GM-CSF cells were a gift from Michael Sykes. L929 cells were obtained from ATCC (CCL-1TM).

### Bone marrow derived macrophage culture and characterization

The animal protocols used for this study were approved by the Institutional Animal Care and Use Committee at MSKCC. For most studies, BMDMs were differentiated from primary mouse bone marrow as previously described(Weischenfeldt and Porse 2008). In brief, femurs from C57BL/6J mice (B6, Jackson #000664) were harvested and bone marrow was extracted by centrifugation. After red blood cell lysis in ACK buffer (10 mM KHCO3, 150 mM NH4Cl, 100 µM EDTA), 3 × 106 bone marrow cells were plated in 10 cm non-TC treated dishes with 10 mL DMEM supplemented with 25 % M-CSF-conditioned media, which was obtained from L929 cells according to the established protocol(Weischenfeldt and Porse 2008). Differentiated BMDMs were transferred to a new plate on day 6 and allowed to attach overnight for experiments on day 7. Day 6 BMDMs were assessed for purity flow cytometrically using antibodies against CD11b, CD11c, and F4/80. For Itgb2 knockout experiments, heterozygous mice were obtained from The Jackson Laboratory (B6.129S7-Itgb2tm2bay/J, #003329) and bred to generate Itgb2+/- and Itgb2-/- breeding pairs. Pups were genotyped by PCR and all experiments used gender-matched littermates for each biological replicate.

### HoxB8ER conditionally immortalized progenitor culture and macrophage differentiation

HoxB8-ER progenitor cells were generated using an established protocol (Wang, 2006 Nature Methods) from bone marrow harvested from B6 (HoxB8-WT) and Cas9 knock-in mice (HoxB8-Cas9) (Jackson Labs #026179). After retroviral transduction of the HoxB8-ER oncogene, cells were cultured in HoxB8 Culture Media (RPMI supplemented with 1 % Pen-Strep, 2 mM L-glutamine, 10 % FBS, 1 µM *β*-estradiol, 5 % GM-CSF conditioned media) for one month to select for clones expressing HoxB8-ER.

To generate CRISPR-KO lines, HoxB8-Cas9 cells were transduced with lentivirus containing targeting or non-targeting sgRNA by spinoculation (1400 × g, 2 h). After spinoculation, the viral media was replaced with HoxB8 Culture Media. After 24 h, half of the media was replaced with fresh HoxB8 Media Culture containing 5 µg/mL blasticidin-HCl (ThermoFisher A1113903) for selection. Cells were kept under selection for one week, with daily exchange into fresh media. CD18 knockout progenitor cells were FACS-sorted after CD18 staining (BioLegend 101414) and recovered for 1 week. Knockout-efficiency was determined by CD18 staining and flow cytometry or by western blot for Talin (Abcam ab157808, Secondary: Goat anti-Mouse 800CW, LI-COR 925-32210) and Vinculin (Abcam ab91459, Secondary: Goat Anti-Rabbit 680RD, LI-COR 926-68071).

To generate f-tractin mCherry reporter lines (HoxB8-f-Tractin), HoxB8-WT cells were transduced with lentivirus containing pCMV-f-Tractin-mCherry, following the same recovery protocol as above, with 5 µg/mL puromycin instead of blasticidin (ThermoFisher A1113803). After 1 week of selection, mCherry-positive cells were sorted by FACS and recovered for an additional week.

To generate HoxB8-ER macrophages, HoxB8-ER progenitor cells were washed three times in 10 mL complete RPMI without *β*-estradiol before transferring to complete BMDM differentiation media (see above), followed by 6 days of culture at 37 °C in non-TC treated dishes (5.3 × 104 cells / cm2). On day 6, cells were trypsinized and replated for downstream experiments.

### Production of iPSC-derived human macrophages

Macrophages were differentiated from hiPSCs using a previously established protocol (Lachmann et al. 2015), with modifications. In brief, hiPSCs were seeded on CF1 mouse embryonic feeders and maintained for 3 days in ESC medium (KO-DMEM (ThermoFisher, 10829-018), 20 % KO-Serum (ThermoFisher, 10828-028), 2 mM L-glutamine (ThermoFisher, 25030-024), 1 % NEAA (ThermoFisher, 11140-035), 0.2 % beta-mercaptoethanol (Thermo Fisher, 31350-010), 1 % Pen/Strep (ThermoFisher, 15140163), with 10 ng/mL bFGF (Peprotech 100-18B), and subsequently for 4 days in ESC media without bFGF. 7 days after seeding, hiPSC colonies were detached in clusters using a 13 min incubation with collagenase type IV (250 IU/mL final concentration) (Thermo Fisher Scientific; 17104019) and transferred to 6-well low adhesion plates in ESC media supplemented with 10 µM ROCK Inhibitor (Sigma; Y0503) to initiate differentiation. The plates were kept on an orbital shaker at 100 rpm for 6 days to allow for spontaneous formation of embryoid bodies (EB) with hematopoietic potential. On D6, 200-500 µm EBs were picked under a dissecting microscope and transferred onto adherent tissue culture plates (2.5 EBs/cm2) for cultivation in Hematopoietic Differentiation (HD) medium (APEL2 (Stem Cell Tech 05270), with 0.5 % Protein free hybridoma medium (ThermoFisher Scientific; 12040077), 1 % pen/strep, 25 ng/mL hIL-3 (Peprotech; 200-03), and 50 ng/mL hM-CSF (Peprotech, 300-25)). Starting from day 18 of the differentiation and then every week onwards for up to two months, macrophage progenitors produced by EBs were collected from suspension, filtered through a 100 µm mesh, plated at a density of 10,000 cells/cm2, and cultivated for 6 days in RPMI1640/GlutaMax (ThermoFisher Scientific; 61870036) medium supplemented with 10 % FBS (EMD Millipore TMS-013-B) and 100 ng/mL human M-CSF before use in downstream experiments. 24 h before the start of an experiment, macrophages were harvested by trypsinization (0.25 % with 1mM EDTA, ThermoFisher 25200056) and plated at a density of 25,000 macrophages per well in an 8-well imaging chamber (Ibidi 80821). Phagocytosis was assessed by live imaging as described below.

### Ex vivo mouse microglia culture

Primary mouse microglia were harvested from P2-P4 B6 mice by first dissociating brains using the Miltenyi gentleMACS Octo Dissociator with Heaters, brain dissociation kits (Miltenyi Biotec 130-096-427, 130-107-677, 130-092-628), and gentleMACS C Tubes (Miltenyi Biotech 130-093-237), following the manufacturer’s instructions. Microglia were purified by magnetic separation using CD11b MicroBeads (Miltenyi Biotec 130-097-142), LS Columns (Miltenyi Biotech 130-042-401) and the QuadroMACS separator (Miltenyi Biotech 130-091-051), following the manufacturer’s instructions. Isolated primary microglia were then cultured in complete DMEM, and phagocytosis was assessed by live imaging in 8-well chambers (Nunc 155409) coated with collagen IV (Corning 354233) at a density of 100,000 cells per well.

### Microparticle synthesis, functionalization, and characterization

Microparticles were synthesized as previously described(Vorselen, Wang et al. 2020, de Jesus, Settle et al. 2023), with modifications. Aqueous acrylamide solutions were prepared containing 150 mM MOPS (pH 7.4), 0.3 % (v/v) tetramethylethylenediamine (TEMED), 150 mM NaOH, and 10 % (v/v) acrylic acid. The mass fraction of total acrylamide was kept constant at 10 % and the cross-linker mass fraction was altered to adjust Young’s Modulus (see Supplemental Table 1). The mixture was sparged with N2 gas for 15 min and kept under N2 pressure. Hydrophobic Shirasu porous glass (SPG) filters (pore size specific to each batch, see Supplemental Table 1) were sonicated under vacuum in n-dodecane, mounted on an internal pressure micro kit extruder (SPG Technology Co), and submerged into the organic phase, consisting of 99 % hexanes and 1 % (v/v) Span-80 (Sigma-Aldrich, S6760), with continuous stirring at 300 rpm. 10 mL of the acrylamide mixture was then loaded into the apparatus and extruded under N2 pressure into 250 mL of the hexane mixture. To initiate polymerization, the resulting emulsion was heated to 60 ºC and 2,2’-azobisisobutyronitrile (2 mg/mL, Sigma Aldrich) was added. The reaction was kept at 60 ºC for 3 hours and then at 40 ºC overnight. Polymerized microparticles were harvested by washing 3 times in hexanes and 2 times in ethanol (30 minutes, 4000 rpm), followed by resuspension in 100 mL phosphate buffered saline (PBS). For some batches, excess ethanol caused incomplete partition of particles into aqueous solution, so additional PBS was added until the particles formed one pellet under centrifugation (4000 rpm, 5 min). Particles were washed two more times in PBS and reduced to an appropriate volume (108 particles / mL). See Supplemental Table 1 for exact extrusion conditions and formulations for each batch of particles used in this study.

For labeling, the particles were streptavidinated using amine coupling as previously described(de Jesus, Settle et al. 2023): briefly, 4.0 × 107 beads were activated with 1-ethyl-3-(3-dimethylaminopropyl)carbodiimide (EDC, Thermo Fisher Scientific [TFS] 22980) and N-hydroxysuccinimide (NHS, TFS 24500) in MES buffer (pH 6.0) with 0.1 % (v/v) Tween 20. Then, the buffer was exchanged into basic PBS (pH 8.0) with 0.1 % Tween-20 for a 2 h conjugation with 10 µM streptavidin (Prozyme, SA10) and 10 µM fluorescent dye (Lissamine Rhodamine B ethylenediamine, FITC ethylenediamine, or Alexa Fluor 488 Cadaverine). Unreacted carboxy-NHS was quenched with excess ethanolamine for 30 min, and the resulting DAAM particles exchanged into PBS (pH 7.4). To functionalize particles for phagocytosis assays, streptavidinated particles were incubated for 2 h in PBS with Biotin-SP-Whole IgG (Jackson Immuno Research 015060003) or Biotin-Phosphatidylserine (Echelon 31B16), followed by 3 PBS washes.

Diameter and Young’s Moduli were measured on particles conjugated to streptavidin, LRB, and FITC, and then coated with Biotin-IgG (10 pmol per million particles, pmol/M). Young’s Modulus was measured by Atomic Force Microscopy (AFM) using large radius hemispherical tips (LRCH-15-750, Team NanoTec). Particles were indented 3 times each and the apparent Young’s modulus was taken as the mean of the 3 measurements. 30 particles were measured for each batch. Particle size was assessed using an inverted fluorescence microscope (Olympus IX-81) fitted with a 20× objective lens (0.75, Olympus). Images of particles (N 39) were taken at the focal plane of maximum diameter, and diameters were subsequently measured using Fiji.

### In vitro phagocytosis of DAAM particles

For flow-cytometry based detection, BMDMs were cultured for 6 days on non-TC treated dishes, trypsinized, and plated in a 24-well non-TC treated plate with 105 cells in 1mL of BMDM differentiation media per well. On the following day, IgG- or PtdS-coated DAAM particles were vortexed and added into each well, followed by centrifugation at 100 × g for 3 min to accelerate particle settling. BMDMs and particles were co-incubated for the specified amount of time (usually 1 h), then washed twice with PBS and removed from the well by trypsinization and scraping. The resulting cells were collected and transferred to a 96-well plate for washing and staining. Cells were washed 3× with 200 µL FACS buffer. In experiments where CD11b or CD18 were measured, cells were stained for 1 h at 4°C in 1:200 conjugated antibody (CD11b-APC (M1/70), Invitrogen 0112-82 or CD18-AF647(M18/2), BioLegend 101414), followed by 2 washes with FACS buffer. Finally, 3 µM DAPI was added 5 min prior to flow cytometric analysis. Measurements were taken on a CytoFLEX benchtop flow cytometer (Beckman Coulter). Live cells were identified using the forward scatter, side scatter, and DAPI channels. Non-conjugate beads were used to define the LRB+FITClo gate. Phagocytosis was measured as LRB+FITCloDAPI-cells divided by the total DAPI-population.

For microscopy-based phagocytosis assays, BMDMs were plated at a density of 2.5 × 104 cells per well in an 8-well chamber slide (Ibidi 80821) and labeled with 1 µg/mL Hoechst 3342 (ThermoFisher H1399). DAAM particles were added and centrifuged at 100 × g for 3 min to encourage contact formation. Imaging was started within 5 min of centrifugation using a ZEISS Axio Observer.Z1 epifluorescence microscope fitted with an X-Cite exacte light source and a 10×/0.45 numerical aperture (NA) objective. 4 channels (FITC, TRITC, DAPI, and Brightfield) and 4 fields of view per well were captured at 2 min intervals for 2 h.

A custom MATLAB script was used to derive phagocytic efficiency and phagocytic index from time-lapse data. Still images were thresholded and segmented by LRB signal to label particles in each field of view. For each labeled particle, mean FITC and LRB intensity values were obtained and the FITC/LRB ratio computed. Every tenth frame (20 min) was processed independently. To compute the threshold for acidified targets, FITC/LRB ratios for all labeled particles across all processed frames were pooled to generate a distribution of represented ratios and the MATLAB functions ksdensity and findpeaks were used to identify two FITChi and FITClo peaks as the two largest peaks in the distribution. The threshold was set as the midpoint between the peaks. In some experiments, the number of phagocytosed particles was low, and only one peak could be identified. In these cases, the threshold was set at 0.5× the peak ratio value. Cells were counted in each well by thresholding and segmenting using the DAPI channel. Phagocytosis efficiency for each time point was calculated as the percentage of labeled particles below the calculated threshold. Phagocytic index was calculated as the number of labeled particles below the threshold divided by the number of cells in the field of view.

### In vivo phagocytosis of DAAM particles

1 mL of aged thioglycolate (TG) was injected intraperitoneally into B6 mice. Following 24 h of elicitation, 107 DAAM particles (coated with 10 pmol/M IgG) were injected intraperitoneally (volume 200 µL in 1× PBS). After 2 h, mice were sacrificed, and cells were isolated by peritoneal lavage as previously described(Ray and Dittel 2010). Samples were then stained with fluorescently conjugated antibodies against CD11b-APC (Invitrogen 0112-82), F4/80-BV421 (BD 565411), Ly6G-BV650 (BD 740554). Peritoneal macrophages were identified as the F4/80+CD11b+Ly6G-cells. Within this population, LRB+FITClo events were identified based on a non-conjugate bead control and phagocytic efficiency was measured as the percentage of CD11b+F4/80+Ly6G-cells in the LRB+FITClo gate.

### In vitro phagocytosis of cancer cells

E0771 cells transduced with doxycycline-inducible MRTF-A or empty vector control were cultured in complete DMEM and grown to 80% confluence in 10 cm TC-treated dishes. 24 hours before experiments, E0771 cells were harvested by trypsinization and cultured at a density of 106 cells/well in 6-well NunclonTM SpheraTM low-adhesion plates (ThermoFisher, 174932) to prevent substrate attachment. 500 ng/mL doxycycline (Sigma, D5207) was then added for 24 h to induce overexpression of MRTF-A. D7 BMDMs stained with CellTraceTM CFSE according to manufacturer’s instructions and were plated in 24-well non TC dishes at 5 × 104 cells per well. On the next day, E0771 target cells were pre-treated with 100 nM MycB or DMSO for 1 h at 37 °C, followed by staining with 1 µM Cypher5e (Sigma GEPA15405) and 1 µg/mL Hoechst 3342 (ThermoFisher H1399) according to manufacturer’s instructions. They were then purified by centrifugation over Histopaque® (Millipore Sigma 10771). 1 × 105 target cells were added to the 24-well BMDM plate and incubated with 1 mg/mL anti-mouse Isotype or anti-CD47 antibodies (BioXCell BE0083 and BE02830). After 2 h, Macrophage-Target conjugates were harvested by trypsinization and analyzed by flow cytometry. Phagocytosis was measured as the ratio of CFSE+Cypher5e+ events over all CFSE+ events. The Cypher5e+ threshold was established using CFSE+Hoechst-events in the isotype control sample.

### Live confocal microscopy and quantification

HoxB8-f-Tractin progenitor cells were differentiated into macrophages as described above and transferred to 8-well chamber slides at a density of 1.5 × 104 cells/ well 24 h prior to the start of the experiment. The chamber-slides were imaged using a SoRa spinning disk confocal microscope (Nikon) fitted with a 63×/1.49 NA objective. To capture images, chamber slides were first placed onto the microscope to focus on the macrophage plane. 50 µL (2.5 × 104) DAAM particles were then pipetted into the center of the well. After 5 min, the region was scanned to identify potential initiating events (particle and cell in contact) and image capture was initiated. Image stacks encompassing the entire particle and macrophage (60-100 slices per stack, 0.3µm slice depth) were captured at the maximum rate obtainable by the system (10-15 s interval per image stack). Time-lapses were captured for a minimum of 10 min, up to 40 min if phagocytosis did not complete. Phagocytic events that did not complete within 30 minutes were considered “stalled.”

To calculate mean phagocytic cup velocity, we first identified the time points at which phagocytosis was initiated (start of contact) and completed (particle fully encompassed in f-tractin). If phagocytosis did not complete, the final timepoint of the movie was used as the end time point. Then for each time point, using Fiji measurement tools, an effective engulfed diameter (d_n) was measured as the distance between the phagocytic cup base and the intersection of the plane of maximum advancement (see Supp Fig 4 for schematic diagram). Instantaneous phagocytic cup velocity was calculated as the distance traveled by actin during the measured period of time:

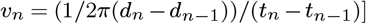

And the mean phagocytic cup velocity was calculated using the total distance traveled during the full engulfment progress:

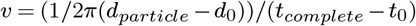

or, if phagocytosis did not complete, the total distance traveled over the entire movie was used:

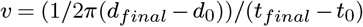

### Fixed confocal microscopy and quantification

To quantify phagocytic cup progress, 5 × 104 macrophages were plated in each well of an 8-well chamber slide (Ibidi 80821), and 105 DAAM particles (IgG-coated, AF488 labeled) were added and centrifuged at 100 × g for 5 min at 4°C to synchronize contact formation. After centrifugation, plates were incubated at 37 °C for 5 min, 10 min, or 30 min and fixed by adding an equal volume of 4 % pre-warmed paraformaldehyde (Electron Microscopy Sciences, final concentration 2 %), followed by 20 min incubation at 37 °C. Each well was then neutralized by removing 200 µL of PFA/Media waste and adding 500 µL complete media (DMEM, 10 % FBS). Neuralization was completed by three washes with 500 µL RPMI per well, followed by 3 washes with 500 µL PBS. Cell-particle conjugates were then permeabilized by 5 min incubation in 0.5 % Triton X-100 in PBS (PBST), followed by a 1 h incubation in PBST with 1 % BSA to block. Samples were then stained for 1 h with Phalloidin-AF594 or Phalloidin-AF568 (1:400 dilution). Following staining, samples were washed 3× with 500 µL PBST and 5× with 500 µL PBS. At each washing step, care was taken to not allow the wells to dry or to disturb particles at the bottom of the well. Samples were imaged using a Leica Stellaris 8 point scanning confocal microscope fitted with a 63×/1.4 NA oil objective. Particle-cell conjugates were identified morphologically by first focusing on particles and then confirming by phalloidin signal that a cell was in contact or nearby. Image stacks were captured using 0.3-µm z-sectioning (80 slices per stack). Phagocytic cup progress was quantified using a custom MATLAB GUI script: each TIFF stack is first presented to the user using the sliceViewer MATLAB function, which prompts the user to click the center of the particle. This center is then used to present an X-Z slice through the center of the particle. The user then outlines the phagocytic cup in the phalloidin channel using the MATLAB drawpolygon function. This enables the particle to be segmented using the particle fluorescence channel and converted into a series of equally spaces points around its circumference. % actin coverage is then calculated as the percentage of particle edge coordinates that fall within the user-defined polygon. To avoid user bias, images from a given experiment were loaded randomly and no identifying information about each image was given to the user. After processing, incidental/non-phagocytic macrophage-particle contacts (<20 % coverage area) were removed from the dataset.

For analysis of CD18-localization, permeabilized cells were first incubated overnight with anti-CD18-AF647 (BioLegend 101414). The following day, they were washed once with PBST, stained with Phalloidin-AF568, processed, and imaged as described above. To analyze F-actin and CD18 accumulation at the phagocytic cup, image stacks were first thresholded and binarized based on AF488 signal to identify the particle volume and boundaries. This mask was then dilated by 4 pixels in all directions to create a “shell” around the particle. Phalloidin and CD18 signal was also thresholded and segmented to identify the cell volume and boundary. Enrichment of each signal at the cup was then calculated as the ratio of the mean fluorescence intensity within the intersection between the particle shell and cell volume divided by the mean fluorescence intensity in the entire cell volume.

To calculate actin and CD18 clearance ratios, we first pre-processed the data using a previously established MATLAB pipeline to render the particles as surface coordinates in 3D cartesian and polar space(Vorselen, Wang et al. 2020, de Jesus, Settle et al. 2023). Phalloidin and CD18 signal intensity was localized to the particles using a 0.3 µm search distance. Fully engulfed particles were removed from this analysis, and particles were rotated such that the base of the phagocytic cup aligned with the bottom of the particle in XYZ coordinates. The rendered particle was transformed to 2D by stereographic projection to a point below the particle in the Z direction (see Figure S4), and two perpendicular line profiles were then measured through the center of the projected interface. To robustly identify the edge of the contact, a custom MATLAB GUI script was used and the peaks of F-actin or CD18 intensity furthest from the contact center in either direction were selected. Clearance ratio was calculated as the mean fluorescence intensity of points in the central 50 % of the contact surface area divided by the mean fluorescence intensity of points in the outer 50 % (25 % in both directions). CD18-F-actin lag distance was calculated as the distance between the phagocytic cup edge as defined by the CD18 signal or the F-actin signal (each identified independently using the aforementioned GUI).

### Phosphoproteomic analysis

Bone marrow from three mice was pooled and differentiated into BMDMs as described above. 24 h before the experiment, BMDMs were trypsinized and replated into 6-well non-TC treated culture plates. IgG-coated (10pmol/M) DAAM particles were overlayed onto the cells, centrifuged for 3 minutes at 300 × g, and then incubated at 37 °C for 10 min. After 10 min, plates were quickly transferred to ice, trypsinized, and snap frozen with liquid nitrogen. One replicate consisted of pooled material from all 6 wells of one plate per stiffness condition. To ensure rapid processing after the addition of DAAM particles, two plates (one with 18kPa particles, one with 0.6kPa) were processed at a time, and all eight replicates were completed in batches over the course of 4 h.

Samples were lysed in buffer containing 8 M urea and 200 mM EPPS (pH at 8.5) with protease inhibitor (Roche) and phosphatase inhibitor cocktails 2 and 3 (Sigma). Benzonase (Millipore) was added to a concentration of 50 U/mL and incubated (RT, 15 min) followed by water bath sonication. Samples were centrifuged at 4 °C, 14,000 × g for 10 min and supernatant extracted. The Pierce bicinchoninic acid (BCA) assay was used to determine protein concentration. Protein disulfide bonds were reduced with 5 mM tris (2-carboxyethyl) phosphine (room temperature, 15 min), then alkylated with 10 mM iodoacetamide (RT, 30 min, dark). The reaction was quenched with 10 mM dithiothreitol (RT, 15 min). Aliquots of 100 ug were taken for each sample and diluted to approximately 100 µL with lysis buffer. Samples were subjected to chloroform/methanol precipitation as previously described1. Pellets were reconstituted in 50 µL of 200mM EPPS buffer and digested with Lys-C (1:50 enzyme-to-protein ratio) at 37 °C for 4 h. Trypsin was then added (1:50 enzyme-to-protein ratio) and the samples incubated overnight at 37 °C.

Anhydrous acetonitrile (ACN) was added to a final concentration of 30%. Peptides were TMT-labeled as described(Navarrete-Perea, Yu et al. 2018). Briefly, peptides were TMT-tagged by adding 10 µL (28 µg/µL) TMTPro reagents (16plex) for each respective sample and incubated for 1 h (RT). A ratio check was performed by taking a 1 µL aliquot from each sample and desalted using the StageTip method (Rappsilber, Mann et al. 2007). TMT-tags were then quenched with hydroxylamine to a final concentration of 0.3% for 15 min (RT). Samples were pooled 1:1 based on the ratio check and vacuum-centrifuged to dryness. Dried peptides were reconstituted in 1 mL of 3% ACN/1% TFA, desalted using a 100 mg tC18 SepPak (Waters), and vacuum-centrifuged overnight.

Phosphopeptides were enriched using the Thermo High-Select Fe-NTA Phosphopeptide Enrichment Kit (Cat. No.: A32992). Unbound peptides and washes were saved and dried down for further analysis of the non-phosphorylated (phospho-depleted) peptides. The phosphopeptide eluate was vacuum-centrifuged to dryness and reconstituted in 100 µL of 1% ACN/25 mM ammonium bicarbonate (ABC). A StageTip was constructed by placing two plugs with a narrow bore syringe of a C18 disk (3M Empore Solid Phase Extraction Disk, #2315) into a 200 µL tip (VWR, Cat. #89079-458). StageTips were conditioned with 100 µL of 100% ACN, 70% ACN/25mM ABC, then 1% ACN/25mM ABC. Phospho-enriched sample was loaded onto the StageTip and eluted into 6 fractions of 3, 5, 8, 10, 12, and 70% ACN/25mM ABC with 100 µL each. Fractions were immediately dried down by vacuum-centrifugation and reconstituted in 0.1% formic acid (FA) for LC-MS/MS. Phospho-depleted peptides were reconstituted in 300 µL 0.1% TFA then fractionated by Pierce High pH Reversed-Phase Peptide Fractionation (Cat. No.: 84868) into 8 fractions (10, 12.5, 15, 17.5, 20, 22.5, 25, and 70% ACN). Fractions were dried by vacuum-centrifugation and reconstituted in 0.1% FA for LC-MS/MS.

Phosphopeptide-enriched and phospho-depleted peptide fractions were analyzed by LC-MS/MS using a Thermo Easy-nLC 1200 (Thermo Fisher Scientific) with a 50 cm (inner diameter 75 µm) EASY-Spray Column (PepMap RSLC, C18, 2 µm, 100 Å) heated to 60 °C and coupled to an Orbitrap Fusion Lumos Tribrid Mass Spectrometer (Thermo Fisher Scientific). Peptides were separated by direct inject at a flow rate of 300 nL/min using a gradient of 5 to 30% acetonitrile (0.1% FA) in water (0.1% FA) over 3 h, then to 50% ACN in 30 min, and analyzed by SPS-MS3. MS1 scans were acquired over a range of m/z 375-1500, 120K resolution, AGC target (standard), and maximum IT of 50 ms. MS2 scans were acquired on MS1 scans of charge 2-7 using an isolation of 0.5 m/z, collision induced dissociation with activation of 32%, turbo scan and max IT of 120 ms. MS3 scans were acquired using specific precursor selection (SPS) of 10 isolation notches, m/z range 110-1000, 50K resolution, AGC target (custom, 200%), HCD activation of 65%, max IT of 150 ms, and dynamic exclusion of 60 s. Raw data files were processed using Proteome Discoverer (PD) version 2.4.1.15 (Thermo Scientific). For each of the TMT experiments, raw files from all fractions were merged and searched with the SEQUEST HT search engine using a mouse UniProt protein database downloaded on 2019/12/13 (92,249 entries). Cysteine carbamidomethylation was specified as a fixed modification, while methionine oxidation, acetylation of the protein N-terminus, TMTpro (K) and /TMTpro (N-term), phosphorylation (STY), and deamidation (NQ) were set as variable modifications. The precursor and fragment mass tolerances were 10 ppm and 0.6 Da respectively. A maximum of two trypsin missed cleavages were permitted. Searches used a reversed sequence decoy strategy to control peptide false discovery rate (FDR) and 1% FDR was set as the threshold for identification. Phosphosite localization was assigned by PD IMP-ptmRS node. Phosphopeptides measured in all replicates were log2-transformed. Unpaired t-test was used to calculate p-values in differential analysis, and volcano plots were generated based on log2FC and p-value.

### Schematic Diagrams created with BioRender.com

The following diagrams were created with BioRender.com: 1A, 1D, 1E, Supp 1D, 2A, 2B, 2C, 2D, 2G, Supp 2A, 3H, Supp 4D, Supp 4E, 4A, 4D, 4E, 4F, Supp 5E, 5A, 5C, 5D, 5E, 5F, 6E, 6F, and 6I.

### Statistics

Graphpad was used for statistical analysis (Graphpad software, Inc). Details can be found in the legend of each figure. N represents number of independent biological replicates.

### Data Availability

Raw proteomic LC/MS data and imaging files are available upon request from the lead author (Morgan Huse).

### Code Availability

All custom MATLAB scripts are available upon request from the lead author (Morgan Huse).

**Supplementary Table 1:** DAAMP Particle Synthesis Formulations and Extrusion Conditions.

| Batch # | Young’s Modulus (kPa) | Diameter Mean (µm) | Mono:Total Acry-lamide Ratio | % Bis-arylamide (by mass) | Membrane Pore Size (µm) | Membrane Lot | Extrusion Pressure (psi) | Span-80 Conc. (v/V) |
| --- | --- | --- | --- | --- | --- | --- | --- | --- |
| 0.31 | 2.54 (±0.4) | 8.31 (±0.8) | 154:1 | 0.065 | 2.4 | PJU15E21 | 3.51 | 1% |
| 0.32 | 1.28 (±0.2) | 9.31 (±1.2) | 250:1 | 0.040 | 2.4 | PJU15E21 | 3.60 | 1% |
| 0.33 | 0.598 (±0.12) | 9.66 (±0.8) | 308:1 | 0.032 | 2.4 | PJU15E21 | 3.42 | 1% |
| 0.34 | 18.0 (±4.1) | 9.65 (±0.8) | 42.7:1 | 0.234 | 3.0 | PJU13A28 | 1.67 | 1% |

